# Optogenetic control of PRC1 reveals that bridging fibers promote chromosome alignment by overlap length-dependent forces

**DOI:** 10.1101/865394

**Authors:** Mihaela Jagrić, Patrik Risteski, Jelena Martinčić, Ana Milas, Iva M. Tolić

## Abstract

During metaphase, chromosome position at the spindle equator is regulated by the forces exerted by kinetochore microtubules and polar ejection forces. However, the role of forces arising from mechanical coupling of sister kinetochore fibers with bridging fibers in chromosome alignment is unknown. Here we develop an optogenetic approach for acute removal of PRC1 to disassemble bridging fibers, and show that they promote chromosome alignment. Tracking of the plus-end protein EB3 revealed longer antiparallel overlaps of bridging microtubules upon PRC1 removal, which was accompanied by misaligned and lagging kinetochores. Kif4A/kinesin-4 and Kif18A/kinesin-8 were found within the bridging fiber and lost upon PRC1 removal, suggesting that these proteins regulate the overlap length of bridging microtubules. We propose that PRC1-mediated crosslinking of bridging microtubules and recruitment of kinesins to the bridging fiber promotes chromosome alignment by overlap length-dependent forces transmitted to the associated kinetochores fibers.

---

Pre-anaphase chromosome movements culminate with chromosome alignment at the spindle equator, a distinctive feature of mitosis important for the synchronous anaphase poleward movement of chromatids and proper telophase nuclear reformation (Fonseca et al., 2019; Maiato et al., 2017). Chromosome congression to the metaphase plate, a process of directed chromosome movement from the polar regions towards the spindle equator, has been explored extensively (Barisic et al., 2014; Cai et al., 2009; Kapoor et al., 2006; Maiato et al., 2017). Yet, the maintenance of chromosome alignment at the spindle equator is less understood. This is a dynamic process given that the chromosomes constantly make small oscillatory excursions across the equator (Skibbens et al., 1993). Two mechanisms have been proposed to underlie the maintenance of chromosome alignment at the equator, namely mechanisms based on length-dependent pulling and polar ejection forces.

Length-dependent pulling forces exerted by kinetochore fibers (k-fibers) are thought to be regulated by kinesin-8 motor proteins, which are needed for proper kinetochore alignment in various organisms from yeast to humans (Garcia et al., 2002; Stumpff et al., 2008; Wargacki et al., 2010). These motors can “measure” microtubule length because they bind to microtubules along their length and move towards the plus end, leading to more kinesin-8 accumulated at the plus end of longer microtubules, where they promote microtubule catastrophe, i.e., a switch from growth to shrinkage (Tischer et al., 2009; Varga et al., 2006). During kinetochore movements across the spindle equator, the leading k-fiber shrinks, while the trailing one grows and accumulates kinesin-8, resulting in microtubule catastrophe followed by the movement of the trailing kinetochore back towards the equator. Although this mechanism can explain kinetochore alignment at the spindle equator (Klemm et al., 2018) the switching dynamics characteristic for this model, where the trailing kinetochore initiates the change of direction of motion, differs from the observations in mammalian cells where the leading kinetochore typically changes the direction before the trailing one (Armond et al., 2015; Civelekoglu-Scholey et al., 2013; Wan et al., 2012).

In addition to the forces produced by k-fibers, polar ejection forces push chromosome arms away from the pole, powered by arm-bound chromokinesins that walk towards the plus end of microtubules (Bajer et al., 1982; Brouhard and Hunt, 2005). Because microtubule density increases towards the pole, these forces help the chromosomes to stay away from the poles, but most likely have little effect on kinetochore movements close to the spindle equator (Cane et al., 2013; Ke et al., 2009). Thus, the current models do not provide a complete picture of kinetochore alignment at the spindle center.

K-fibers are surrounded by a dense network of spindle microtubules with which they have multiple interactions (McDonald et al., 1992; O’Toole et al., 2020), resulting in forces acting on k-fibers and thus also on kinetochores. In particular, each pair of sister k-fibers is tightly linked by the bridging fiber, a bundle of antiparallel microtubules that balances the tension on sister kinetochores (Kajtez et al., 2016; Polak et al., 2017; Vukusic et al., 2017). However, the role of forces exerted by bridging fiber in chromosome alignment at the metaphase plate is unknown.

In metaphase, overlap regions within bridging fibers are crosslinked by protein regulator of cytokinesis 1 (PRC1) (Kajtez et al., 2016; Polak et al., 2017; Tolic, 2018). PRC1, like other non-motor microtubule-associated proteins from Ase1/PRC1/MAP65 family, selectively bundles antiparallel microtubules and provides stable overlaps *in vitro* (Bieling et al., 2010b; Janson et al., 2007; Mollinari et al., 2002; Subramanian et al., 2010). Cellular studies of its function show that PRC1 is associated with the spindle midzone in anaphase, where its activity is essential for stable microtubule organization, localization of numerous microtubule-associated proteins within this structure, and successful completion of cytokinesis, while its microtubule-binding and -bundling affinities are regulated by phosphorylation and dephosphorylation events (Gruneberg et al., 2006; Jiang et al., 1998; Kurasawa et al., 2004; Liu et al., 2009; Mollinari et al., 2002; Mollinari et al., 2005; Neef et al., 2007; Subramanian et al., 2013; Subramanian et al., 2010; Zhu and Jiang, 2005; Zhu et al., 2006).

In this work, we developed an optogenetic approach for acute and reversible removal of PRC1 from the spindle to the cell membrane, building upon ideas of dimerization or dissociation induced chemically (Cheeseman et al., 2013; Haruki et al., 2008; Robinson et al., 2010; Wordeman et al., 2016) or by light (Fielmich et al., 2018; Guntas et al., 2015; Okumura et al., 2018; van Haren et al., 2018; Yang et al., 2013; Zhang et al., 2017) to rapidly redistribute proteins. By using our assay on metaphase spindles, we found that bridging fibers promote kinetochore alignment. PRC1 removal resulted in partial disassembly of bridging fibers and elongation of their overlap zones. Moreover, the metaphase plate widened, inter-kinetochore distance decreased, and lagging chromosomes appeared more frequently, showing that PRC1 indirectly regulates forces acting on kinetochores. Kif4A/kinesin-4 and Kif18A/kinesin-8 were found to localize in the bridging fiber during metaphase, and were lost upon PRC1 removal. These results, together with the finding that Kif4A or Kif18A depletion by siRNA led to elongated overlaps, suggest that these proteins regulate the overlap length of bridging microtubules. PRC1 removal did not affect the localization of Kif4A on the chromosomes and CLASP1, Kif18A, and CENP-E/kinesin-7 on the plus ends of k-fibers, arguing against perturbed polar ejection forces or molecular events at the kinetochore microtubule plus-ends as origins of the observed kinetochore misalignment. In conclusion, our optogenetic experiments show that bridging microtubules buffer chromosome movements, thus promoting their alignment. We propose that this occurs via length-dependent forces, which depend of the antiparallel overlap length within the bridging fiber.

## RESULTS

### Optogenetic system for fast and reversible removal of PRC1 from the metaphase spindle

To study the role of PRC1 and the forces arising from coupling of bridging and k-fibers in chromosome alignment, we developed an optogenetic tool for fast and reversible removal of PRC1 from the spindle to the cell membrane, based on the previously designed improved light inducible dimer (iLID) system (Guntas et al., 2015). We attached PRC1 to the red fluorescent protein tgRFPt and the bacterial protein SspB, while the iLID, which contains the bacterial peptide SsrA and the light-oxygen-voltage (LOV2) domain, is bound to the cell membrane by a short peptide, CAAX. In this system, LOV2 adopts a conformation that allows dimerization of SsrA and SspB upon exposure to the blue light (**Fig. 1 A**). After cessation of exposure to the blue light, LOV2 adopts its initial conformation leading to decreased affinity of SsrA to SspB. Therefore, exposure to the blue light should induce translocation of PRC1 from the central region of the metaphase spindle, which we will refer to as the spindle midzone, to the cell membrane, whereas cessation of exposure to blue light should restore PRC1 localization on the spindle (**Fig. 1 A**).

**Figure 1.**
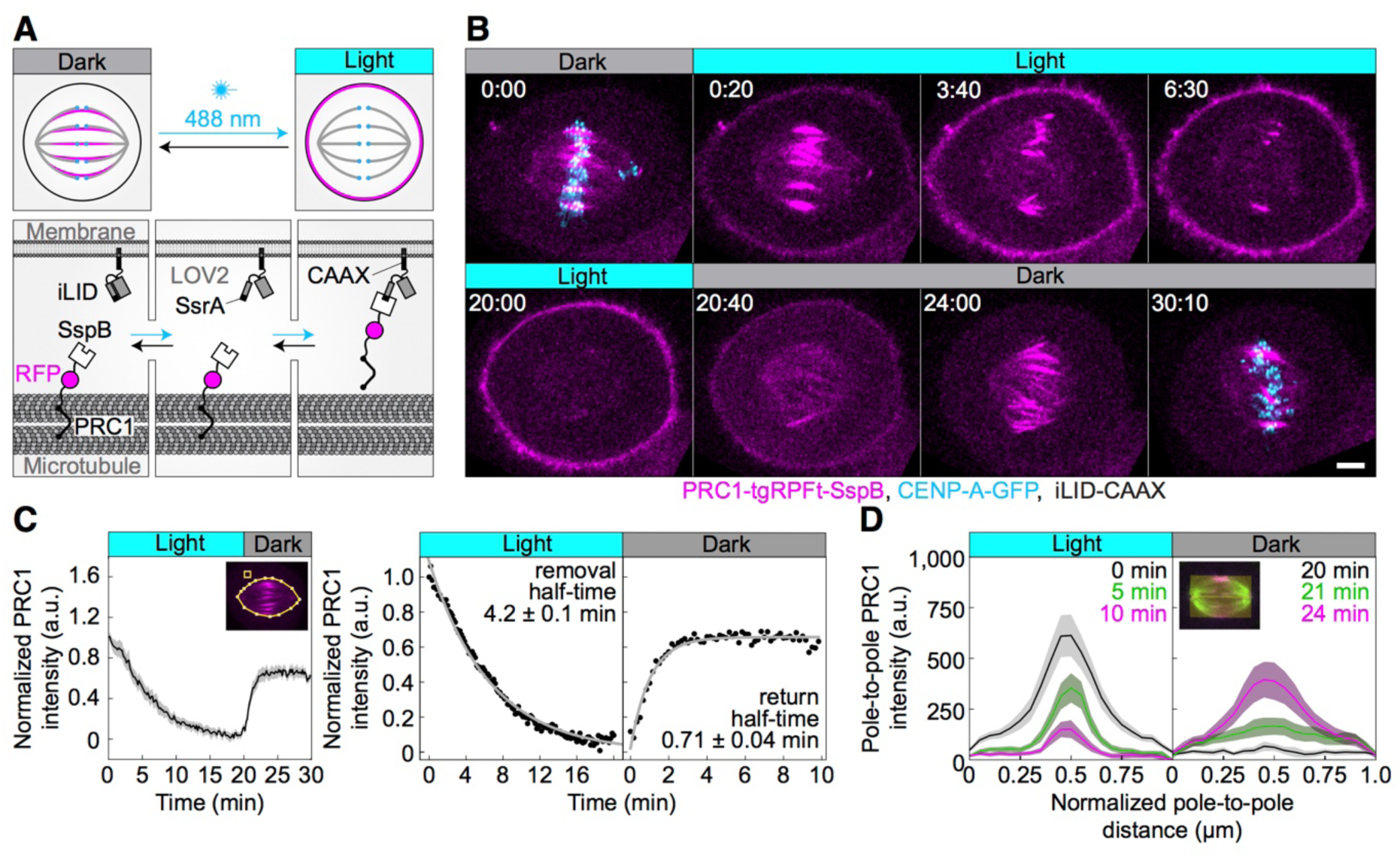
Optogenetic reversible removal of PRC1 from the spindle in metaphase. (**A**) Schematic representation of the optogenetic system. PRC1 is fused with SspB and tgRFPt (opto-PRC1, see **Methods**). iLID, composed of photosensitive LOV2 domain and SsrA is tagged with CAAX sequence which mediates its binding to the cell membrane. Exposure to the blue light induces conformational change in LOV2 domain, enabling dimerization of SspB and SsrA, and thus translocation of PRC1 from the spindle to the cell membrane. After the blue light is turned off, LOV2 adopts its initial conformation, leading to decreased affinity of SspB for SsrA, and consequently dissociation of PRC1 from the membrane and its return to the spindle. (**B)** Time-lapse images of a metaphase spindle in a U2OS cell stably expressing CENP-A-GFP (cyan), depleted for endogenous PRC1, with transient expression of opto-PRC1 (magenta) and iLID-CAAX. Note that kinetochores are shown only in the first and the last time frame in order to better visualize PRC1 removal. Images are maximum intensity projections of three z-planes, smoothed with 0.1-µm-sigma Gaussian blur. Time; min:sec. Scale bar; 5 µm. (**C)** Normalized intensity of opto-PRC1 signal on the spindle (left panel) during its removal (0-20 min) and return (20-30 min). N=15 (see **Fig. S1 F** for individual cells). Scheme depicts the areas where opto-PRC1 intensity was measured: spindle (large polygon) and cytoplasm (small square). Exponential fit (gray lines in the right panel) on mean normalized opto-PRC1 spindle intensity (black points) during 20 min of removal and 10 min of return. Formulae y=A*exp(-τ*x) and y=A*exp(-τ*x)+c were used for opto-PRC1 removal and return, respectively. Parameters for PRC1 removal: A=1.111, τ=0.00277 s^-1^ (RSE=0.03), and return: A=-0.635, c=0.656, τ=0.01622 s^-1^ (RSE=0.03). The half-time was calculated by ln2/τ. (**D**) Pole-to-pole opto-PRC1 intensity during removal (left; N=14) and return (right; N=9) to the spindle. Mean and s.e.m are color-coded corresponding to the time when measured (upper right corners). Scheme depicts the area where opto-PRC1 intensity was measured (yellow) to obtain the mean intensity projection onto the pole-to-pole axis. Mean (thick lines); s.e.m. (shaded areas); N (number of cells).

To test our optogenetic approach, we used U2OS cells with stable expression of CENP-A-GFP, transient expression of PRC1-tgRFPt-SspB (henceforth opto-PRC1) and iLID-CAAX (henceforth opto cells; **Fig. 1 B**; **Video 1**). Endogenous PRC1 was depleted 90 ± 2% (all results are mean ± s.e.m.) by siRNA before addition of opto-PRC1 (**Fig. S1 A**). Before exposure to the blue light, opto-PRC1 had normal localization on the microtubule bundles in the spindle midzone (**Fig. 1 B**; 0:00 min), with the length of PRC1 streaks of 3.77 ± 0.08 µm (n=193 bundles, N=30 cells), consistent with that of endogenous and fluorescently labeled PRC1 in metaphase (Kajtez et al., 2016; Polak et al., 2017), though the total signal intensity of opto-PRC1 on the spindle was higher compared to endogenous PRC1 (**Fig. S1 B**). Addition of opto-PRC1 did not change the duration of metaphase, as inferred from the fraction of cells that entered anaphase during image acquisition, which was similar in cells with endogenous PRC1 and cells treated with PRC1 siRNA and opto-PRC1 (79 ± 6 % N=37, and 71 ± 5 % N=72, respectively; p=0.4, Pearson’s Chi-squared test; **Fig. S1 C**). After exposure to the blue light, opto cells were able to progress to cytokinesis (**Fig. S1 D**). Taken together, these data suggest that opto-PRC1 replaces the depleted endogenous PRC1.

Upon exposure to the blue light, opto-PRC1 signal on the spindle decreased and its signal on the membrane increased (**Fig. 1 B**; 0:20-20:00 min). After the blue light was switched off, opto-PRC1 returned to the spindle midzone (**Fig. 1 B**; 20:40-30:10 min). In control experiments without the blue light or without iLID (henceforth control), opto-PRC1 was not removed (**Fig. S1 E**). Thus, our optogenetic approach allows for acute and reversible control of PRC1 localization in metaphase.

To quantify the dynamics and spatial pattern of opto-PRC1 removal and return, we measured the intensity of opto-PRC1 on the metaphase spindle (**Fig. S1 F,G**). We found that 88 ± 3% of opto-PRC1 was removed after 20 min of exposure to the blue light with a half-time of 4.2 ± 0.1 min (**Fig. 1 C**). During the opto-PRC1 removal, there was simultaneous decrease in both signal intensity and length of the overlap region (**Fig. 1 D**, left; **Fig. S1 G**, top), which may be due to fewer antiparallel regions being positioned laterally than in the central part of the bundle (Mastronarde et al., 1993). The signal of the outermost midzone bundles typically lasted longer than of the inner ones (**Fig. 1 B**; 3:40**-**6:30). After the blue light was switched off, opto-PRC1 signal restored to 65 ± 1% of the initial intensity within 10 minutes, with return half-time being 0.71 ± 0.04 min (**Fig. 1 C**). The faster PRC1 return to the spindle in comparison with its removal may be due to the higher affinity difference between PRC1 binding to the spindle and to the membrane in the dark than under light. During the opto-PRC1 return, it initially localized throughout the spindle, with gradual increase in intensity in the spindle midzone (**Fig. 1 D**, right; **Fig. S1 G**, bottom), suggesting that PRC1 has higher unbinding rate outside than within the overlap bundles in the spindle. This result is consistent with PRC1 having a life-time of several seconds on single microtubules and a 10-fold preference for overlap regions *in vitro* (Subramanian et al., 2010).

### Acute PRC1 removal during metaphase leads to misaligned kinetochores

To test whether bridging fiber has a role in the maintenance of chromosome alignment, acute optogenetic removal of PRC1, a depletion of which is known to perturb bridging fibers (Polak et al., 2017) should affect chromosome positioning at the spindle equator (**Fig. 2 A**). Surprisingly, we observed that the acute removal of opto-PRC1 resulted in movement of sister kinetochore pairs away from the metaphase plate (**Fig. 2 B; Video 2**), which is not found after long-term PRC1 depletion by siRNA (Polak et al., 2017). Upon opto-PRC1 removal, the distances of sister kinetochore midpoints from the equatorial plane increased (d_EQ_, **Fig. 2 B,C,D**; **Fig. S2 A,B**). While >95% of kinetochore pairs were found within the region of PRC1 streaks, i.e., less than 2 µm away from the equatorial plane before removal of opto-PRC1, 9.3 ± 2.7% of kinetochore pairs made excursions far outside this region after opto-PRC1 removal (**Fig. S2 B**). The displaced kinetochores were found more often in the inner part of the spindle, i.e., close to the spindle axis, than in the outer regions (**Fig. 2 E**). Kinetochores fluctuated to a similar extent in the presence and absence of opto-PRC1, but in its absence the displaced kinetochores fluctuated within a region that was offset from the equatorial plane (**Fig. S2 D**). These displaced kinetochores upon opto-PRC1 removal had lower inter-kinetochore distance in comparison to non-displaced ones, suggesting that kinetochore displacement was related to a more severe reduction of tension (**Fig. 2 F**). On average, kinetochores remained displaced even after opto-PRC1 return (**Fig. 2 D**). We conclude that PRC1 has a role in keeping kinetochores in tight alignment on the metaphase plate.

**Figure 2.**
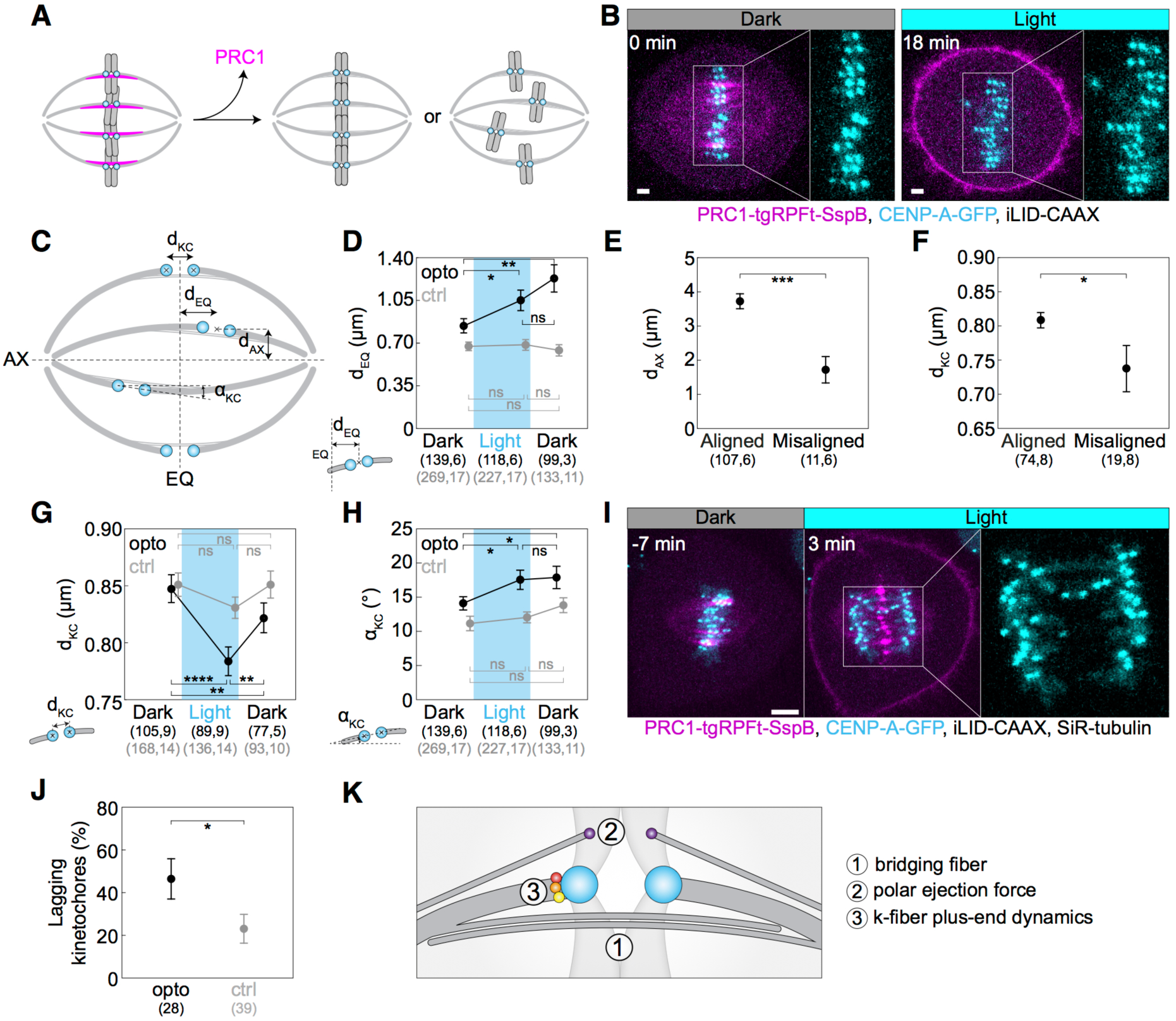
Optogenetic removal of PRC1 disrupts kinetochore alignment on the metaphase plate and leads to lagging kinetochores in anaphase. **(A)** Schematic representation of possible outcomes of acute removal of PRC1 from the spindle on chromosome alignment. (B) Spindle in a U2OS cell stably expressing CENP-A-GFP (cyan) with transient expression of opto-PRC1 (magenta) and iLID-CAAX before (0 min, Dark), and at the end of continuous exposure to the blue light (18 min, right). Enlargements show kinetochores only. Scale bar; 2 µm. (C) Schematic of measured parameters. d_KC_, inter-kinetochore distance; d_EQ_, distance between sister kinetochore midpoint and equatorial plane (EQ); α_KC_, angle between sister kinetochore axis and spindle long axis (AX); d_AX_, distance between sister kinetochore midpoint and spindle long axis. (D) Measurements of d_EQ_ in opto (black) and control (gray) cells before (0 min, Dark), at the end of continuous exposure (20 min, Light) and 10 min after cessation of exposure to the blue light (30 min, Dark), in U2OS cells expressing CENP-A-GFP. (E) d_AX_ of aligned (d_EQ_<2 µm) and misaligned (d_EQ_=2.5 ± 0.2 µm) kinetochore pairs upon PRC1 removal. (F) d_KC_ of aligned and misaligned kinetochore pairs upon PRC1 removal. (G) Measurements of d_KC_. Legend as in D. (H) Measurements of α_KC_. Legend as in D. (I) Time-lapse images of a spindle in a U2OS cell as in B, stained with SiR-tubulin (not shown). Anaphase onset is at time 0 min. Lagging kinetochores can be seen at 3 min (middle). Enlargement shows kinetochores only. (J) Occurrence of lagging kinetochores in anaphase of opto (black) and control (gray) U2OS cells. (K) Schematic of three mechanisms that could be involved in kinetochore alignment. Cyan rectangles inside graphs indicate exposure to the blue light. Numbers in brackets denote measurements and cells; single numbers denote cells. In D, G, H opto cells are without SiR-tubulin, and control cells include those with and without SiR-tubulin. All images are maximum intensity projections of three z-planes, smoothed with 0.5-pixel-sigma Gaussian blur. Error bars; s.e.m. Scale bar; 5 µm. Statistical analysis; t-test (D-H), two-proportions z-test (J). p-value legend: < 0.0001 (****), 0.0001 to 0.001 (***), 0.001 to 0.01 (**), 0.01 to 0.05 (*), ≥ 0.05 (ns).

The mean inter-kinetochore distance (d_KC_, **Fig. 2 C**) was reduced when opto-PRC1 was removed (**Fig. 2 G**; **Fig. S2 C**), and the distance after 20 min of exposure to the blue light (0.79 ± 0.01 µm) was closer to metaphase (0.87 ± 0.01 µm) than prometaphase (0.66 ± 0.01 µm) values (see **Methods**), suggesting that tension was not completely lost and that these changes were not due to kinetochore detachment from k-fibers (**Fig. S2 C**). In agreement with this, the fraction of cells that entered anaphase during image acquisition was similar in control and opto cells (71 ± 5 % N=72, and 60 ± 5 % N=93, respectively; p=0.1, Pearson’s Chi-squared test; **Fig. S1 C**), indicating that PRC1 removal did not prevent spindle assembly checkpoint satisfaction. After opto-PRC1 return to the spindle, inter-kinetochore distance increased, implying restoration of tension, though not to the original value (**Fig. 2 G**).

To investigate the influence of acute PRC1 removal on the orientation of sister kinetochores, we measured the angle between sister kinetochore axis and long spindle axis (α_KC_, **Fig. 2 C**). We observed that removal of opto-PRC1 caused misorientation of sister kinetochores, i.e., increased α_KC_ (**Fig. 2 B,H**; **Fig. S2 A,E**). Misoriented kinetochores were found at a larger distance from the equatorial plane and closer to the long spindle axis (**Fig. S2 F**). Interestingly, sister kinetochore pairs remained misoriented even after opto-PRC1 return (**Fig. 2 H**). Similarly, PRC1 bundles were misoriented upon PRC1 return (see **Methods**, **Fig. S2 G**). These results suggest that when PRC1 returns to the overlaps whose geometry was perturbed by PRC1 removal, it likely confines the chromosomes in new positions and orientations.

The observed effects of PRC1 removal on inter-kinetochore distance, kinetochore alignment and orientation did not change when SiR-tubulin was added (compare **Fig. 2 D,G,H** without SiR-tubulin and **Fig. S2 B,C,E** including SiR-tubulin stained cells). The effects of PRC1 removal were found neither in control experiments without iLID, nor in a different set of control experiments where cells without opto-PRC1 and without iLID were exposed to the same laser illumination protocol (**Fig. 2 D,G,H**; **Fig. S2, A-C, E**), suggesting that the observed effects were not a consequence of laser photodamage (Douthwright and Sluder, 2017). We conclude that PRC1 plays a role in maintaining kinetochore alignment and orientation within the metaphase plate.

### PRC1 removal during metaphase increases the frequency of lagging kinetochores in anaphase

To test to what extent the acute removal of PRC1 during metaphase affects chromosome segregation, we measured the frequency of lagging kinetochores. We found that lagging kinetochores occurred more frequently when opto-PRC1 was being removed than in control cells (**Fig. 2 I,J**; **Video 3**). Opto cells that showed lagging kinetochores in anaphase had a slightly smaller inter-kinetochore distance before anaphase than opto cells without lagging kinetochores (**Fig. S2 H**), suggesting that a decrease in tension may be involved in the imperfect kinetochore segregation. The cells with lagging kinetochores did not have a larger average kinetochore misalignment in metaphase (**Fig. S2 H**), which indicates that misalignment and lagging kinetochores are not linked on the cell level, though they may be linked locally on individual kinetochores.

As perturbation of the PRC1-CLASP1 interaction and the consequent absence of CLASP1 from the spindle midzone results in lagging chromosomes (Liu et al., 2009), we tested CLASP1 and found that it did not accumulate between segregating chromosomes in opto HeLa cells stably expressing EGFP-CLASP1 (**Fig. S2 I**). Thus, the observed higher occurrence of lagging kinetochores could be attributed to changes in tension during metaphase, perturbed recruitment of CLASP1 to the spindle midzone by PRC1 during early anaphase, or a combination of both effects.

### Acute PRC1 removal leads to longer antiparallel overlap zones within the bridging fibers

Factors that could contribute to the altered chromosome alignment and occurrence of lagging kinetochores upon opto-PRC1 removal are changes related to 1) microtubules in the bridging fibers, 2) polar ejection forces, and/or 3) proteins that modulate the dynamics of k-fiber plus-ends (**Fig. 2 K**). Because PRC1 crosslinks microtubules within bridging fibers (Kajtez et al., 2016; Polak et al., 2017), we first tested the effects of PRC1 removal on those microtubules.

An important aspect of the bridging fiber that may affect chromosome alignment is the dynamics of microtubules that make up these fibers. To explore their dynamics, we developed an assay to track the growing plus ends of individual microtubules in the bridging fiber by using cells expressing the plus end marker EB3 (Stepanova et al., 2003) tagged with GFP (**Fig. 3**). We followed single EB3 spots in the spindle and identified the ones belonging to a bridging fiber as the spots that move towards a kinetochore, cross the region between this kinetochore and its sister, and move beyond it towards the other spindle pole (**Fig. 3 A-C; Video 4**). We found 1.8 ± 0.2 EB3 spots per minute per bridging fiber in control cells, showing that bridging fibers are dynamically remodeled during metaphase (**Fig. 3 D**). This number was similar after opto-PRC1 removal (**Fig. 3 D**), suggesting that the number of dynamic microtubules in the bridge is largely independent of PRC1.

**Fig. 3.**
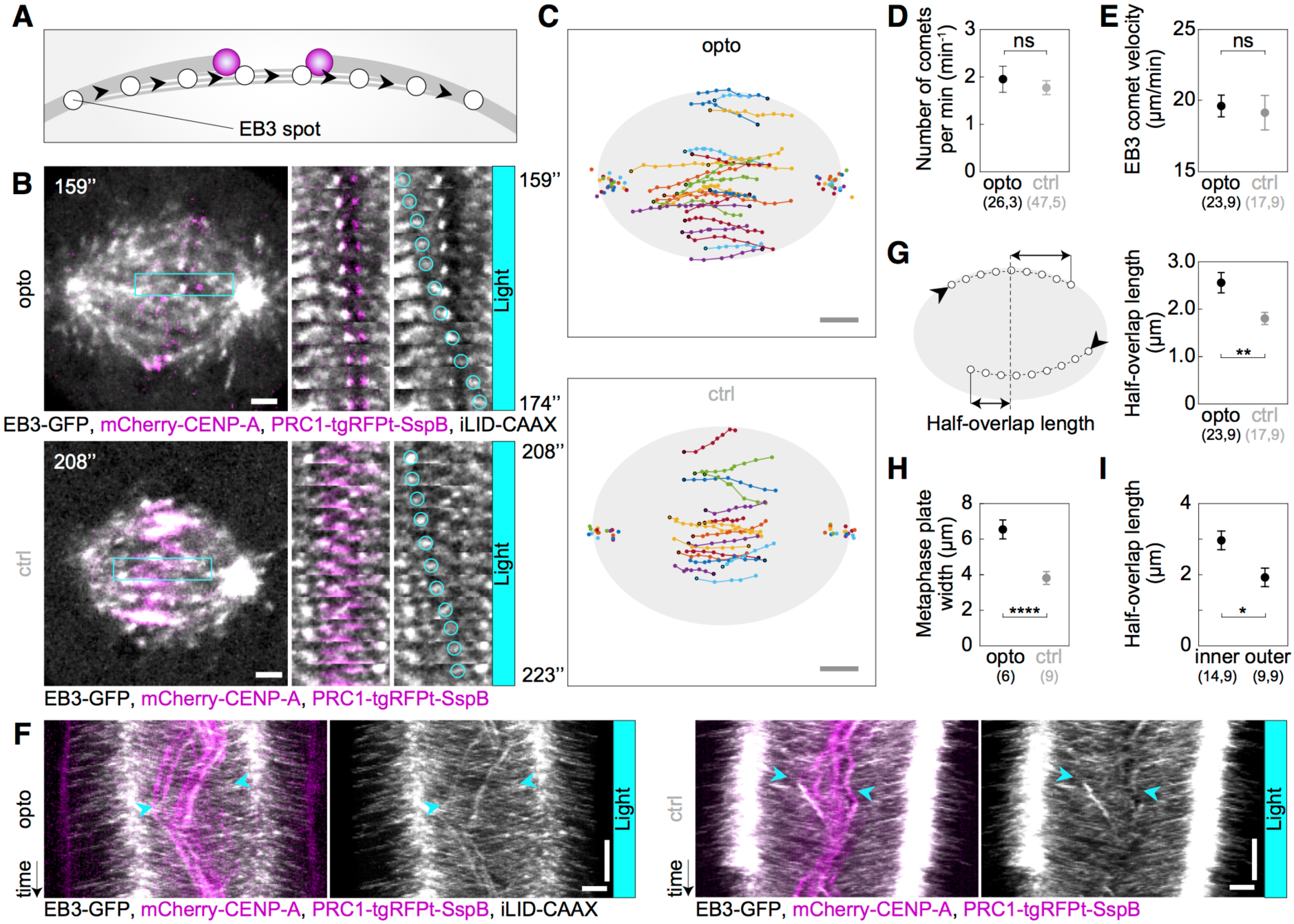
Acute removal of PRC1 elongates antiparallel overlaps within the bridging fibers. (**A**) Schematic of the trajectory of an EB3-marked plus end (white circles) within the bridging fiber, defined as a spot passing the sister kinetochore region (magenta). (**B**) Spindles (left) in U2OS cells with stable expression of 2xGFP-EB3 (gray) and mCherry-CENP-A (magenta), depleted for endogenous PRC1, with transient expression of opto-PRC1 and iLID-CAAX (opto; top) and opto-PRC1 only (control; bottom). Montage of the boxed region over time is shown as merged channels (middle) and GFP (right; the tracked spot is encircled). Time after 10 min of imaging protocol required for removal of PRC1 is shown. Images are single z-planes smoothed with 0.5-pixel-sigma Gaussian blur. (**C**) Trajectories of tracked EB3 spots (connected dots) in opto (top) and control (bottom) cells. Black dot; start of trajectory. Single dots on the sides; spindle poles. (**D**) Number of EB3 spots per minute within the bridging fiber in opto (black) and control (gray) cells. (**E**) EB3 spot velocity within the bridging fiber in opto (black) and control (gray) cells. (**F**) Kymographs of opto (left) and control (right) cells after 10 min of imaging protocol required for removal of PRC1, merge (left) and GFP (right). Cyan arrowheads mark the beginning and end of an individual EB3 spot trajectory. Note the difference in the position of track end with respect to the equatorial plane. (**G**) Half-overlap length (left) defined as the distance (double arrow) between the end-point of the EB3 spot trajectory and the equatorial plane (dashed line). Black arrowhead; start of trajectory. Half-overlap length in opto (black) and control (gray) cells measured as in **A** and retrieved from tracks shown in **B** (right). (**H**) Metaphase plate width in opto (black) and control (gray) cells measured from kymographs as the largest distance between kinetochore pairs positioned on the opposite sides of the spindle equator in the first two minutes after 10 min of imaging protocol required for removal of PRC1. In control cells, the metaphase plate width corresponds to PRC1-streak length. (**I**) Half-overlap length in opto cells for inner (d_AX_≤2 µm) and outer (d_AX_>2 µm) overlaps. Filled cyan rectangles indicate exposure to the blue light. Numbers in brackets denote measurements and cells; single numbers denote cells. Error bars; s.e.m. Scale bars; 2 µm. Statistical analysis; t-test. p-value legend as in **Fig. 2**.

To assess the changes in the dynamics of bridging microtubules, we followed EB3 spots in the bridge from the time when they can be distinguished from neighboring spots near the pole until they disappear in the opposite spindle half, which we interpret as the moment when the microtubule stops growing (Maurer et al., 2012). Interestingly, EB3 tracks were longer after opto-PRC1 removal than in control cells (**Fig. 3 B,C; Video 4**), but the velocities of the EB3 spots were not affected by opto-PRC1 removal (**Fig. 3 E**). In agreement with these results, kymographs of the central region of the spindle show that EB3 spots reach deeper into the opposite half of the spindle after opto-PRC1 removal (**Fig. 3 F; Fig. S3 A**). Thus, acute removal of PRC1 results in longer bridging microtubules, which is not a consequence of altered microtubule growth rate, but most likely due to a reduced microtubule catastrophe rate.

EB3 tracks allowed us to estimate the length of the overlap zone of antiparallel microtubules in the bridging fiber, which currently cannot be measured after PRC1 removal because PRC1 itself is the only available marker of antiparallel overlaps in the spindle. We define the overlap half-length as the distance beyond the equatorial plane that an EB3 spot covers while moving within the bridging fiber from one spindle half into the other, where this microtubule can form antiparallel overlaps with the oppositely oriented microtubules extending from the other pole (**Fig. 3 G, left**). In control cell, the overlap half-length was 1.8 ± 0.2 µm (n=17 spots in N=9 cells; **Fig. 3 G, right**), which corresponds well to the overlap half-length measured as the half-length of PRC1 streaks in this cell line, 0.5x(3.8 ± 0.4) µm (N=9 cells; **Fig. 3 H**), validating our method for overlap length measurement based on EB3 tracks. After opto-PRC1 removal, the overlap half-length increased to 2.6 ± 0.2 µm (n=23 spots in N=9 cells; **Fig. 3 G**). In contrast, treatment with PRC1 siRNA did not result in a change of overlap length with respect to mock siRNA (**Fig. S3, B-E**). Thus, the antiparallel overlaps became on average 40% longer after acute, but unchanged after long-time PRC1 removal.

Interestingly, the overlaps were especially long in the inner part of the spindle, whereas those on the spindle periphery were similar to overlaps in control cells (**Fig. 3 I**). This spatial difference is correlated with our findings that PRC1 is removed faster from the inner bundles and that displaced and misoriented kinetochores are found more often in the inner than in the outer spindle region, suggesting a mechanistic link between the overlap length and kinetochore positioning.

### Removal of PRC1 reduces the number of microtubules in the bridging fibers

To test to what extent PRC1 removal in metaphase affects the number of microtubules in the bridging fibers (**Fig. 4 A**), we visualized microtubules by using SiR-tubulin, a far-red tubulin dye excited by red light (Lukinavicius et al., 2014), which allowed us to observe microtubules both when the blue light is turned on and switched off. Intensity profiles across the spindle midzone revealed that SiR-tubulin intensity maxima were lower upon PRC1 removal and increased after its return (**Fig. S4 A**; **Video 5).** Measurements of SiR-tubulin signal intensity between and lateral from sister kinetochores showed that upon PRC1 removal the tubulin signal was reduced specifically in the bridging fibers (**Fig. 4, A-C**; **Video 5**). As an alternative to SiR-tubulin, which is a taxol-based dye that may affect microtubule dynamics (Lukinavicius et al., 2014), we tested YFP-tubulin, but the excitation laser for YFP also activated the optogenetic system (Wang and Hahn, 2016) (**Fig. S4 B**).

**Figure 4.**
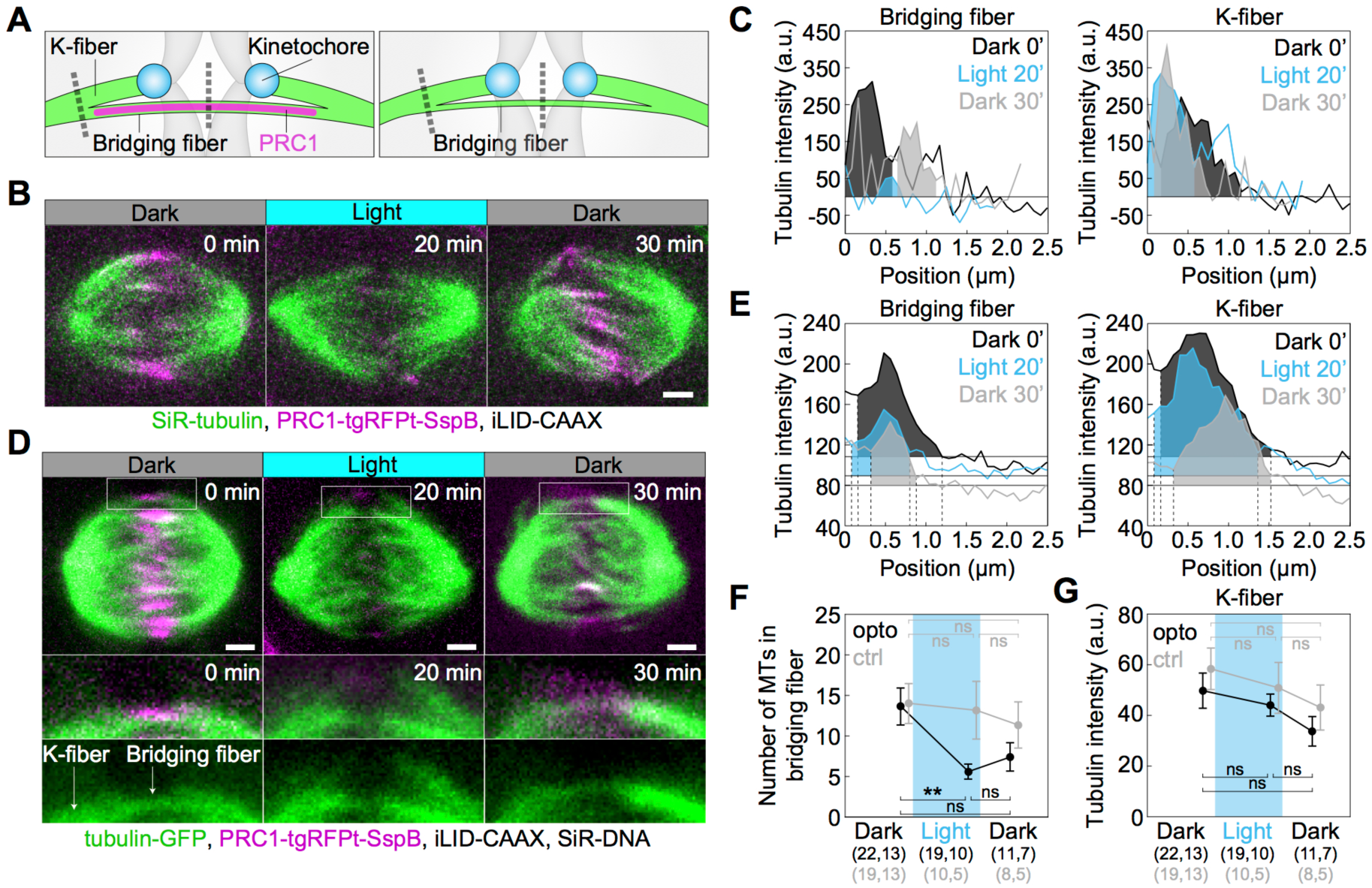
Optogenetic removal of PRC1 reduces bridging fibers and straightens the spindle contour. (**A**) Schematic of PRC1 (magenta) removal from bridging fibers and positions where bridging fiber and k-fiber intensities were measured (dashed lines). (**B**) Spindle in a U2OS cell with stable expression of CENP-A-GFP (not shown), depleted for endogenous PRC1, with transient expression of opto-PRC1 (magenta) and iLID-CAAX, and stained with SiR-tubulin (green), before exposure to the blue light (0 min, Dark), at the end of continuous exposure to the blue light (20 min, Light), and 10 min after cessation of exposure to the blue light (30 min, Dark). Images are a single z-plane smoothed with 0.5-pixel-sigma Gaussian blur. (**C**) Background-corrected SiR-tubulin intensity profiles of the bridging fiber (left) and k-fiber for cell shown in **B** (0 min, black; 20 min, cyan; 30 min, gray). (**D**) Spindle in a HeLa cell with stable expression of tubulin-GFP (green), depleted for endogenous PRC1, with transient expression of opto-PRC1 (magenta) and iLID-CAAX, and stained with SiR-DNA (not shown), before exposure to the blue light (0 min, Dark, left), at the end of continuous exposure to the blue light (20 min, Light, middle) and 10 min after cessation of exposure to the blue light (30 min, Light, right). Enlargements of the boxed region (middle: merge, bottom: GFP) are shown. Note that at 20 min opto-PRC1 is absent from the spindle. (**E**) Graphs showing tubulin-GFP intensity profiles of the bridging fiber (left) and k-fiber (0 min, black; 20 min, cyan; 30 min, gray) for cell shown in **D**. Horizontal line marks the background signal, vertical dashed lines delimit the area (shaded) where signal was measured. (**F**) Number of microtubules in the bridging fiber in opto HeLa cells (i.e., where PRC1 was removed; black) and control (gray) in same time-points as in **D**. The bridging fiber intensity in control cells before exposure to the blue light is set to correspond to 14 microtubules (see **Fig. S4 D)**. (**G**) Tubulin-GFP signal intensity laterally from k-fiber tip in opto (black) and control (gray) at time-points as in **D**. Cyan rectangles inside graphs indicate exposure to the blue light. Numbers in brackets; number of measurements and cells, respectively. Error bars; s.e.m. Scale bars; 2 µm. Statistical analysis; t-test. p-value legend as in **Fig. 2**.

Finally, we used tubulin-GFP to determine tubulin signal intensities of the bridging fibers and k-fiber tips upon acute removal of PRC1 (**Fig. 4 A,D,E**; **Fig. S4 C-F**; see **Methods**). Upon exposure to the blue light, tubulin signal intensity in the bridging fibers decreased ∼2.5 fold, which corresponds to 5.6 ± 0.9 microtubules given that the average number of microtubules in the bridging fiber is 14 (Kajtez et al., 2016). Together with the finding that the number of growing microtubules in the bridging fiber is similar with and without PRC1 (**Fig. 3 D**), this result implies that in the presence of PRC1 the bridge contains more microtubules with a smaller fraction of them being dynamic than without PRC1, and that PRC1 removal leads mainly to disassembly of non-dynamic microtubules. Upon PRC1 return the intensity and thus the number of microtubules remained low (**Fig. 4, D-F**; **Fig. S4 D**), possibly as a consequence of the perturbed spindle architecture in the absence of PRC1, which may be able to recover after a longer time period. Importantly, the intensity at the position lateral from the k-fiber tips was unaltered (**Fig. 4 G**), suggesting that the k-fibers were not affected by PRC1 removal. Thus, acute removal of PRC1 and its return change the number of microtubules specifically in the bridging fibers.

Given that the bridging fiber is under compression (Kajtez et al., 2016), reduction of number of microtubules in the bridging fiber is expected to reduce this compression and thus to straighten the k-fibers (**Fig. S4 G**). Tracking of the pole-to-pole contour of the outermost k-fibers revealed that PRC1removal indeed straightened the k-fibers and thus made the spindle diamond-shaped (**Fig. S4, H-K**). This result provides further support to the finding that the bridging fibers largely disassemble upon acute PRC1 removal.

Since bridging fibers are nevertheless present upon PRC1 removal, we asked how the residual microtubules are bundled together. Eg5/kinesin-5, which localizes in the bridging fibers (Kajtez et al., 2016; Mann and Wadsworth, 2018), was still detectable in these fibers after PRC1 siRNA (**Fig. S4 L**). Thus, we propose that microtubule crosslinkers such as Eg5 crosslink the remaining microtubules in the bridge after acute PRC1 removal.

### Kif4A, Kif18A, and MKLP1 localize in the bridge during metaphase in a PRC1-dependent manner

To investigate the mechanism of bridging microtubule regulation via PRC1, we analyzed the localization of major proteins that regulate spindle microtubule dynamics and/or are binding partners of PRC1: Kif4A, CLASP1, Kif18A, and CENP-E, and MKLP1 (Al-Bassam et al., 2010; Bieling et al., 2010b; Bringmann et al., 2004; Gruneberg et al., 2006; Kurasawa et al., 2004; Liu et al., 2019; Maiato et al., 2003; Mayr et al., 2007; Stumpff et al., 2008; Stumpff et al., 2012; Yu et al., 2016) before and after PRC1 removal in metaphase (**Fig. 5** and **Table 1**).

**Figure 5.**
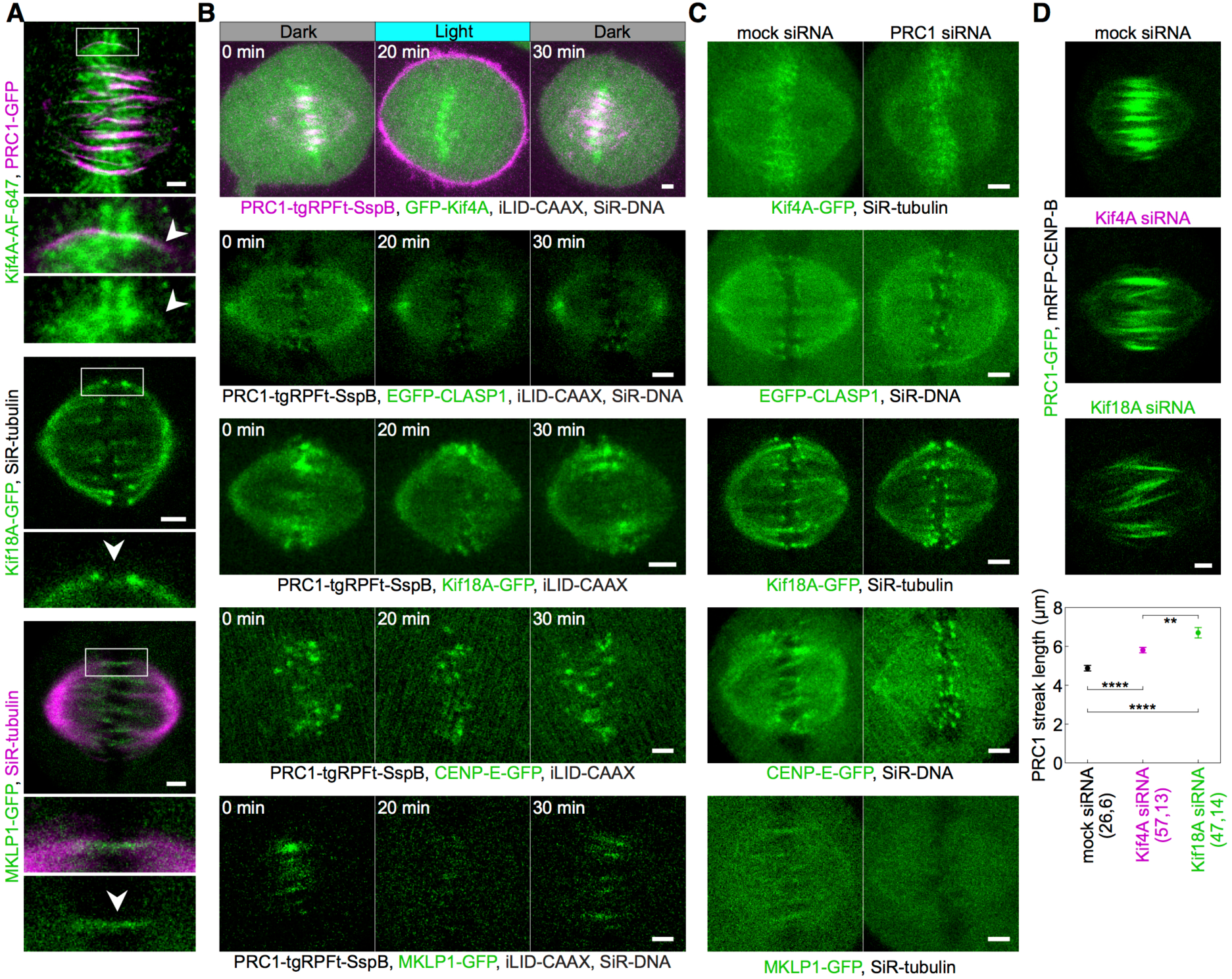
Localization of Kif4A, CLASP1, Kif18A, CENP-E and MKLP1 after acute removal and long-term depletion of PRC1. (**A**) Spindle (top block) in a HeLa cell stably expressing PRC1-GFP (magenta) immunostained for Kif4A-AF-647 (green). Arrows in the enlargements of the boxed region point to Kif4A outside chromosomes, at the positions where PRC1-GFP is found. Spindle (middle block) in a HeLa cell stably expressing Kif18A-GFP (green) and stained for SiR-tubulin (not shown) and spindle (bottom block) in a HeLa cells stably expressing MKLP1-GFP (green) and stained for SiR-tubulin (magenta). Arrows in the enlargements of the boxed regions point to the bridging fiber. (**B**) Time-lapse images of: unlabeled U2OS cell with transient expression of opto-PRC1 (magenta), iLID-CAAX and GFP-Kif4A (green), and stained with SiR-DNA (not shown) (first row); HeLa cell with stable expression of EGFP-CLASP1 (green) with transient expression of opto-PRC1 and iLID-CAAX, and stained with SiR-DNA (second row); unlabeled U2OS cell with transient expression of opto-PRC1, iLID-CAAX and EGFP-Kif18A (green) (third row); unlabeled U2OS cell with transient expression of opto-PRC1, iLID-CAAX and GFP-CENP-E (green) (fourth row); HeLa BAC cell stably expressing MKLP1-GFP (green) and with transient expression of opto-PRC1 and iLID-CAAX, and stained with SiR-DNA (fifth row). All time points are before (0 min, Dark), at the end of continuous exposure (20 min, Light) and 10 min after cessation of exposure to the blue light (30 min, Dark). (**C**) Images of mock (left column) and PRC1 siRNA treated (right column): HeLa BAC cell expressing Kif4A-GFP (green) stained with SiR-tubulin (first row); HeLa cell stably expressing EGFP-CLASP1 (green) stained with SiR-DNA (second row); HeLa BAC cell expressing Kif18A-GFP (green) (third row); HeLa BAC cell expressing CENP-E-GFP (green) stained with SiR-DNA (fourth row); HeLa BAC cell expressing MKLP1-GFP (green) stained with SiR-tubulin (fifth row). (**D**) Spindles from HeLa cells stably expressing PRC1-GFP (green), treated with mock (top), Kif4A (middle), and Kif18A (bottom) siRNA. Graph shows the lengths of PRC1 streaks per treatment. Numbers in brackets; number of measurements and cells, respectively. Images in **A** (bottom), **B**, and **C** are maximum intensity projections of 2-3 z-planes, smoothed with 0.5-pixel-sigma Gaussian blur. Scale bars; 2 µm.

**Table 1.** Localization of proteins in the bridging fiber and at kinetochores after acute and long-term removal of PRC1. For long-term depletion, p-values for localization in the bridging fiber - Kif4A: 0.043; MKLP1: 8*10^-6^; CLASP1: 0.113; Kif18A: 0.033. Statistics: Pearson’s Chi-squared test. In acute removal protein is considered present if at least 83% of cells had it in the bridge and absent if it was undetectable in at least 66% of inspected cells. Numbers in brackets; number of cells. Legend: + present, - absent, +/- unclear. *in the case of Kif4A kinetochore (KC) localization includes chromosome arms.

|  | Acute removal |  |  |  | Long-term depletion |  |  |  |
| --- | --- | --- | --- | --- | --- | --- | --- | --- |
|  | Dark (0 min) |  | Light (20 min) |  | mock |  | siRNA |  |
|  | Bridge | KC | Bridge | KC | Bridge | KC | Bridge | KC |
| Kif4A* | + | + | +/- | + | + | + | - | + |
|  | (6) | (6) | (6) | (6) | (10) | (10) | (16) | (16) |
| CLASP-1 | +/- | + | +/- | + | + | + | + | + |
|  | (5) | (5) | (5) | (5) | (15) | (15) | (8) | (8) |
| Kif18A | + | + | - | + | + | + | - | + |
|  | (6) | (6) | (6) | (6) | (18) | (18) | (8) | (8) |
| CENP-E | +/- | + | +/- | + | +/- | + | +/- | + |
|  | (5) | (5) | (5) | (5) | (6) | (6) | (10) | (10) |
| MKLP-1 | + | - | - | - | + | - | - | - |
|  | (6) | (6) | (6) | (6) | (19) | (19) | (9) | (9) |

We first looked at the localization of these proteins in the bridging fiber. Surprisingly, we found that Kif4A, Kif18A, and MKLP1 localize in the bridge during metaphase, visible as thin lines across or next to the location of sister kinetochores (**Fig. 5 A**). These proteins were not detectable in the bridge after both acute (opto) and long-term (siRNA) PRC1 removal, and returned to the bridge upon PRC1 return (**Fig. 5 B,C,** see also **Fig. S5 A,B for Kif18A; Table 1**). CLASP1 and CENP-E also showed weak localization in the bridge, which in the case of CENP-E agrees with a recent study that clearly showed its presence in the bridge by using super-resolution microscopy (Steblyanko et al., 2020).

Among the tested proteins, only MKLP1 was exclusively localized in the bridging fibers, co-localizing with PRC1 (**Fig. 5, A-C**; **Fig. S5, C-F**). Even though MKLP1 was completely removed from the spindle by acute PRC1 removal, it was not detected at the cell membrane together with PRC1. It may be that MKLP1 binds rather weakly to PRC1 in metaphase and/or that the absence of PRC1 decreases its affinity for microtubules. In addition, the ability of MKLP1 to bind along scaffold of antiparallel overlaps could depend on the role of PRC1 in dictating 35-nm-inter-microtubule spacing, proposed to be important to enable localization of specific proteins within these structures (Kellogg et al., 2016; Subramanian et al., 2010).

Kif4A and Kif18A are known to regulate microtubule length (Mayr et al., 2007; Varga et al., 2006; Wandke et al., 2012). Thus, we hypothesized that these kinesins may also regulate the length of bridging microtubules and hence their antiparallel overlaps during metaphase. Indeed, removal of Kif4A or Kif18A by siRNA resulted in longer PRC1-labeled overlaps (**Fig. 5 D**). Given that Kif4A and MKLP1 are involved in microtubule sliding within the bridging fiber in anaphase (Vukušić et al., 2019), our results suggest that Kif4A, Kif18A, and MKLP1 regulate bridging microtubule dynamics and/or sliding in metaphase, thereby controlling the geometry of their overlaps.

### Kif4A localization on chromosome arms and CLASP1, Kif18A, and CENP-E on kinetochore fiber tips does not depend on PRC1

As an alternative to the changes in the bridging fiber, disruption of polar ejection forces may be the cause of kinetochore misalignment upon PRC1 removal (**Fig. 2 K**), in particular because these forces are modulated by Kif4A (Stumpff et al., 2012). The most prominent Kif4A localization in metaphase is on chromosomes (Gruneberg et al., 2006; Kurasawa et al., 2004; Zhu and Jiang, 2005) (**Fig. 5 B**; **Fig. S5 G,H**). Neither acute nor long-term PRC1 removal had an effect on the Kif4A signal on the chromosomes (**Fig. 5 B,C**; **Fig. S5 I; Table 1**). Thus, Kif4A localization on chromosomes does not depend on PRC1, suggesting that the observed changes in kinetochore alignment are not caused by disruption of polar ejection forces. In early anaphase, the amount of Kif4A on segregated chromosomes was similar in opto and control cells (**Fig. S5 J)**, indicating that increased occurrence of lagging kinetochores was neither due to perturbed polar ejection forces in that phase nor defects in chromosome architecture/condensation.

Finally, kinetochore misalignment and lagging kinetochores upon PRC1 removal may be due to disrupted localization of proteins that regulate microtubule dynamics at the plus-end of the k-fiber (**Fig. 2 K**). To investigate this possibility, we analyzed the k-fiber localization of the key proteins with this function, CLASP1, Kif18A, and CENP-E (Al-Bassam et al., 2010; Maffini et al., 2009; Maiato et al., 2003; Mayr et al., 2007; Stumpff et al., 2008; Stumpff et al., 2012; Yu et al., 2016). These proteins were localized at the plus-ends of k-fibers before and after acute and long-term removal of PRC1 (**Fig. 5 B,C**; **Table 1**), indicating that kinetochore misalignment upon PRC1 removal are not caused by simultaneous removal of proteins that regulate microtubule dynamics at the k-fiber plus ends.

### Acute and long-term removal of PRC1 result in partially different effect on spindle microtubules and kinetochores

Previous reports have shown that acute rapamycin-dependent protein translocation and long-term depletion by siRNA can yield different and even opposite phenotypes (Cheeseman et al., 2013; Wordeman et al., 2016). To explore to what extent acute optogenetic removal of PRC1 affects the spindle differently than long-term depletion by siRNA, we compared the phenotypes obtained by these two approaches (**Table 2**). Similarly as acute removal, long-term depletion of PRC1 decreased the number of microtubules in the bridging fiber and caused straightening of outermost k-fibers, whereas spindle length and width was unchanged (**Fig. S6, A-D**). However, the effects of acute removal of PRC1 were more severe than those of siRNA (**Table 2**). The two methods decreased the inter-kinetochore distance to a similar extent (**Table 2**; **Fig. S6 E**), even though unlike acute removal, long-term removal reduced the fraction of cells that entered anaphase (35 ± 8 % N=37, and 79 ± 6 % N=37, for PRC1 siRNA treated and untreated, respectively; p=0.046, Pearson’s Chi-squared test; **Fig. S1 C**). Strikingly, in contrast to acute removal, the long-term depletion did not cause kinetochore misalignment or misorientation (**Fig. S6 F,G**). Moreover, only acute removal resulted in longer antiparallel overlaps within bridging fibers (**Fig. 3G; Fig. S3E**). Finally, both long-term and acute PRC1 reduction increased the frequency of lagging kinetochores during early anaphase (**Fig. S6 H**).

**Table 2.** Comparison of effects of acute optogenetic removal of PRC1 and long-term depletion by siRNA. All values are given as mean ± s.e.m. The numbers in the brackets denote the number of measurements and cells, respectively; a single number is the number of cells. Symbols (arrows and equal signs) denote trend of change of parameters; equal sign means no change; longer arrow marks stronger effect. *calculated from our previous study (Polak et al., 2017) ^#^consistent with our previous studies (Kajtez et al., 2016; Polak et al., 2017)

| Parameter | Acute removal |  |  |  | Long-term depletion |  |  |  |
| --- | --- | --- | --- | --- | --- | --- | --- | --- |
|  | 0 min | 20 min | p-value |  | untreated | siRNA | p-value |  |
| **N**<br>**(microtubules)**<br>**in the**<br>**bridging fiber** | 14 ± 2<br>(22,13) | 5.6 ± 0.9<br>(19,10) | 0.0038 | ↓ | 14 ± 1<br>(16, 9)* | 10 ± 1<br>(21,6)* | 0.048 | ↓ |
| **Curvature**<br>($\mu\text{m}^{-1}$) | 0.134 ± 0.004<br>(40, 10) | 0.081 ± 0.005<br>(39, 10) | 6*10<sup>-11</sup> | ↓ | 0.137 ± 0.006<br>(20, 5) | 0.108 ± 0.004<br>(52,13) | 0.0014 | ↓ |
| **θ (°)** | 142 ± 3<br>(20, 10) | 125 ± 3<br>(20, 10) | 2*10<sup>-4</sup> | ↓ | 145 ± 3<br>(10, 5) | 137 ± 2<br>(26, 13) | 0.051 | = |
| **d<sub>KC</sub> (μm)** | 0.87 ± 0.01<br>(226,18) | 0.79 ± 0.01<br>(186, 18) | 1*10<sup>-8</sup> | ↓ | 0.85 ± 0.01<br>(75, 8)<sup>#</sup> | 0.780 ± 0.008<br>(202, 17)<sup>#</sup> | 4*10<sup>-6</sup> | ↓ |
| **d<sub>EQ</sub> (μm)** | 0.78 ± 0.04<br>(290, 17) | 0.94 ± 0.05<br>(240, 16) | 0.035 | ↑ | 0.69 ± 0.05<br>(107, 8) | 0.69 ± 0.03<br>(333, 17) | 0.64 | = |
| **d<sub>EQ</sub>>2 μm (%)** | 4.1 ± 1.2<br>(290, 17) | 8.3 ± 1.8<br>(240, 16) | 0.043 | ↑ | 3.3 ± 1.0<br>(107, 8) | 0.9 ± 0.9<br>(333, 17) | 0.33 | = |
| **α<sub>KC</sub> (°)** | 13.8 ± 0.7<br>(290, 17) | 18.4 ± 1.0<br>(240, 16) | 0.0013 | ↑ | 13.3 ± 1.0<br>(107, 8) | 10.3 ± 0.7<br>(333, 17) | 1*10<sup>-4</sup> | ↓ |
| **α<sub>KC</sub>>35° (%)** | 6.2 ± 1.4<br>(290, 17) | 12.1 ± 2.1<br>(240, 16) | 0.018 | ↑ | 4.7 ± 2.0<br>(107, 8) | 3.6 ± 1.0<br>(333, 17) | 0.83 | = |
| **Lagging**<br>**kinetochores**<br>(%) | 23 ± 7<br>(39) control | 46 ± 9<br>(28) opto | 0.044 | ↑ | 5 ± 4<br>(37) | 40 ± 11<br>(20) | 0.0036 | ↑ |

## DISCUSSION

### An optogenetic system for acute, selective, and reversible removal of spindle proteins

We developed an optogenetic approach that offers acute light-controlled removal of proteins from a normally formed spindle at a precise phase of mitosis. The main advantages of this approach over chemically-induced protein translocation (Cheeseman et al., 2013; Haruki et al., 2008; Robinson et al., 2010; Wordeman et al., 2016) are its reversibility, allowing the protein to return to its initial location within about a minute, and applicability to individual cells. Unlike previous optogenetic approaches (Fielmich et al., 2018; Okumura et al., 2018; van Haren et al., 2018; Yang et al., 2013; Zhang et al., 2017), this method allows for global loss-of-function of full-length spindle proteins, relying on simple protein tagging rather than domain splitting, with no need of chromophore addition. Moreover, this method may be implemented with other optical perturbations (Milas et al., 2018) and used as “*in vivo* pull-down” for probing protein-protein interactions in different phases of the cell cycle. However, this approach depends on high turnover of the protein in comparison with the time scale of interest.

We found differences in phenotypes upon acute PRC1 removal from the spindle by optogenetics and the long-term PRC1 depletion by siRNA. These differences are hard to explain by different levels of PRC1 on the spindle as both methods decreased PRC1 by ∼90%. It is also unlikely that the differences are caused by the interaction of the membrane-translocated PRC1 with astral microtubules because of the uniform PRC1 signal on the membrane, fast return to the spindle, and no change in spindle positioning. Therefore, the generally weaker effects of siRNA in comparison with the acute optogenetic removal are most likely due to compensatory mechanisms being triggered during long-term depletion.

### A model for chromosome alignment by overlap length-dependent forces within the bridging fibers

By overcoming temporal limitations of siRNA, our work reveals an unexpected role of PRC1 and bridging fibers in the regulation of chromosome alignment on the metaphase spindle via overlap length-dependent forces (**Fig. 6A**). We propose that the interactions between k-fibers and bridging fibers regulate the movement of bi-oriented chromosomes along the pole-to-pole axis. If a kinetochore pair is displaced towards one spindle pole, more motors and/or crosslinkers accumulate in the overlap facing the opposite pole because this overlap is longer, and pull the kinetochores back to the center (**Fig. 6 B**). As the efficiency of centering depends on the relative asymmetry in the overlap length on either side, shorter overlap length leads to more precise centering of the kinetochores (**Fig. 6 A,B**).

**Figure 6.**
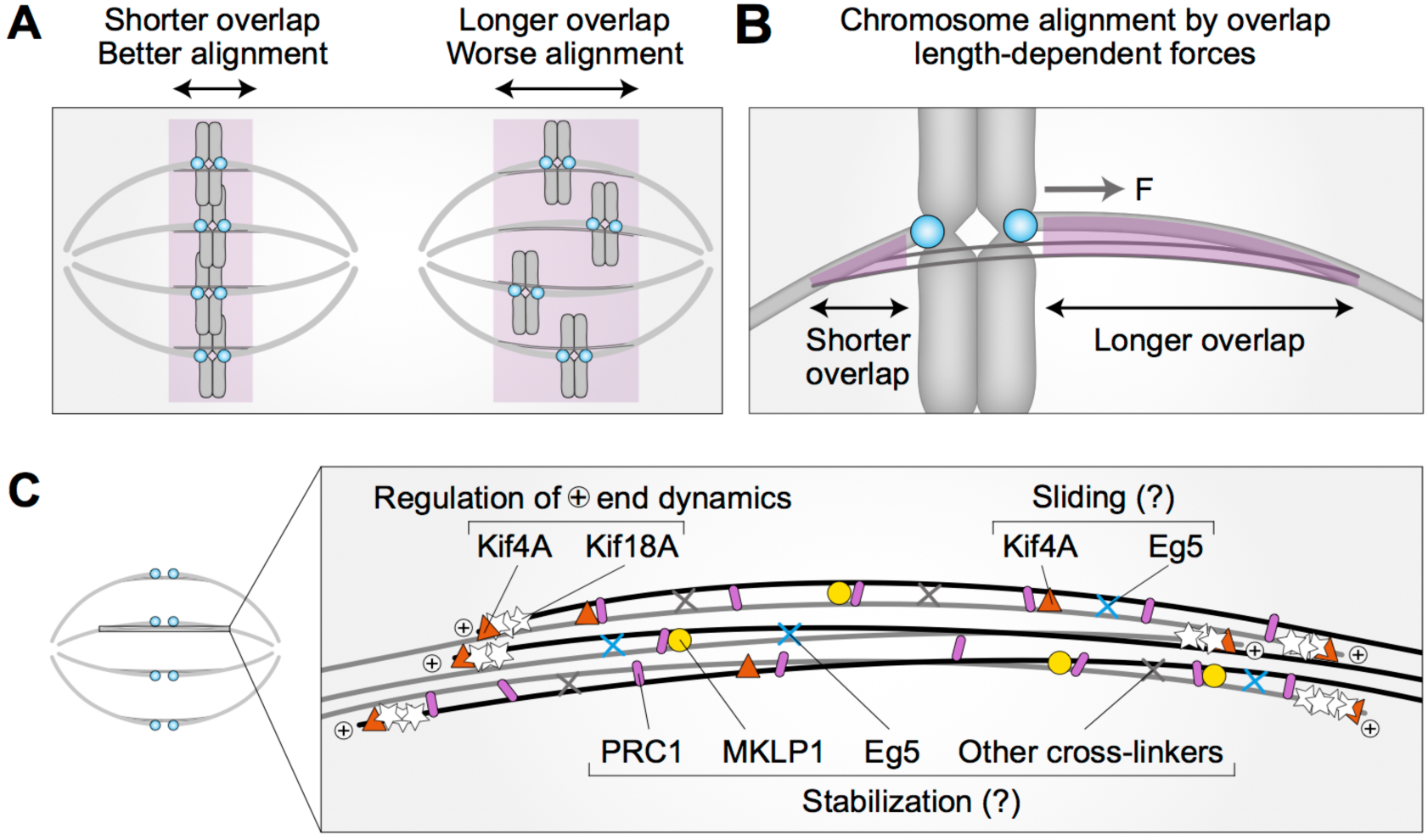
Model for chromosome alignment by overlap length-dependent forces within the bridging fiber. (**A**) We propose that the interactions between k-fibers and bridging fibers regulate the movement of bi-oriented chromosomes by forces that depend on the length of the antiparallel overlaps (purple). Shorter overlaps lead to more precise alignment of kinetochores (cyan) than longer ones. (**B**) If a kinetochore pair is displaced away from the equatorial plane towards one pole, the overlap between the k-fiber and the bridging fiber (purple) is shorter on this side and longer on the opposite side. More motors and/or crosslinkers accumulate in the longer overlap, pulling the kinetochore back to the center (*F*, pulling force). The efficiency of centering depends on the relative asymmetry in the overlap length on either side. This asymmetry is larger if the overlap is short, which explains why short overlaps lead to better alignment than long ones (A). (**C**) The overlap length is regulated by Kif4A and Kif18A at the plus ends of bridging microtubules. Kif4A and Eg5 within the bridging fiber possibly slide the microtubules apart, whereas PRC1 stabilizes the overlaps probably together with MKLP1, Eg5, and other crosslinkers.

How is the length of the bridging microtubules and their overlaps controlled? Interestingly, we found that Kif4A, Kif18A, and MKLP1 localize in the bridging fibers in metaphase and this localization was lost after optogenetic or siRNA-mediated PRC1 removal. As both Kif4A and Kif18A promote microtubule depolymerization (Bieling et al., 2010b; Bringmann et al., 2004; Mayr et al., 2007; Stumpff et al., 2008; Stumpff et al., 2012; Varga et al., 2006), and our experiments showed that depletion of either of them by siRNA leads to longer PRC1-labeled overlaps, the loss of these kinesins from the bridge upon acute PRC1 removal may underlie the observed increase in length of the bridging microtubules. Even though it was reported that Kif4A does not directly interact with PRC1 before late mitosis (Gruneberg et al., 2006; Kurasawa et al., 2004; Zhu and Jiang, 2005), there may be a small pool of Kif4A that does so and binds to the antiparallel overlaps of bridging microtubules in metaphase. PRC1 removal likely results in the removal of this pool of Kif4A from the bridge, which may lead to excessive microtubule polymerization and thus longer overlap zones in metaphase, similar to the long overlaps in anaphase after Kif4A depletion (Hu et al., 2011). Kif18A, which is known to accumulate at microtubule plus-ends where it promotes depolymerization (Gupta et al., 2006; Mayr et al., 2007; Stumpff et al., 2008; Varga et al., 2006), likely regulates also bridging microtubule length in this manner. However, as Kif18A is not a binding partner of PRC1, it is possible that it remains in the bridge after PRC1 removal, but its signal may be undetectable due to the smaller number of microtubules in the bridge and the fact that their plus ends are positioned away from the equator, thus overlapping with the Kif18A signal along the k-fiber. We propose that both Kif4A and Kif18A regulate the length of bridging microtubules and their overlaps (**Fig. 6 C**), with Kif4A likely being more accountable for overlap elongation upon acute PRC1 removal.

The proteins that we found in the bridging fiber, Kif4A, Kif18A, and MKLP1, may also slide apart and/or stabilize the bridging microtubules (**Fig. 6 C**). We also observed Eg5 in the bridging fiber (Kajtez et al., 2016; Mann and Wadsworth, 2018), though its localization was independent of PRC1 in contrast to the other kinesins that we tested. During early anaphase, Kif4A and Eg5 drive the sliding of antiparallel microtubules that elongates the spindle (Vukušić et al., 2019). Thus, these kinesins may have a similar role during metaphase. This possibility is in agreement with previous work showing that Kif4A depletion reduces microtubule poleward flux in metaphase (Wandke et al., 2012). Similarly, Kif18A in the bridging fiber may have microtubule sliding and crosslinking activities equivalent to those of the yeast kinesin-8 (Su et al., 2013). Furthermore, MKLP1 contributes to the stabilization of bridging microtubules in early anaphase (Vukusic et al., 2017), and we suggest that it may perform a similar function in metaphase together with other motors and crosslinkers including Eg5 and PRC1. The roles of these and other proteins within bridging fibers in the regulation of microtubule dynamics and sliding will be an intriguing topic for future studies.

The localization of Kif4A on chromosome arms, where it is involved in polar ejection forces (Bieling et al., 2010a; Brouhard and Hunt, 2005) was unaffected by acute PRC1 removal. Similarly, CLASP1, Kif18A, and CENP-E remained on plus-ends of k-fibers. Although we cannot exclude potential subtle changes in the localization of these proteins or mislocalization of other proteins, our results suggest that the perturbed kinetochores alignment is unlikely due to reduced polar ejection forces or perturbed k-fiber plus ends.

The changes upon acute PRC1 removal were not spatially uniform across the spindle. The most affected part was the inner part of the spindle, where the PRC1 signal disappeared faster and the bridging microtubules became longer than on the periphery of the spindle. The inner bridging fibers were more severely affected by PRC1 removal possibly because they are made up of fewer microtubules than the outer bridges. Severely misaligned kinetochores that moved more than 2 µm away from the equatorial plane and lagging kinetochores occurred also more often in the inner part of the spindle. This local effect is in line with weak mechanical coupling between neighboring k-fibers yet strong coupling between sister k-fibers (Elting et al., 2017; Suresh et al., 2020; Vladimirou et al., 2013), and indicative of a mechanistic link between the bridging fiber geometry and kinetochore alignment.

In conclusion, we propose that overlap length-dependent forces help to position the chromosomes at the equatorial plane of the spindle. The overlap length is regulated by Kif4A and Kif18A within the bridging fiber. Kif4A, Eg5, and possibly other kinesins may slide bridging microtubules apart, whereas PRC1 together with the kinesins stabilizes the overlaps. Thus, in addition to the forces generated at k-fiber tips and polar ejection forces, proper chromosome alignment requires forces generated within the bridging fiber, which are transferred to the k-fiber and rely on precise regulation of the overlap region.

## Methods

### Cell lines

Experiments were performed using: unlabeled human osteosarcoma U2OS cell line and U2OS cell line stably expressing CENP-A-GFP, used in our previous work (Vukusic et al., 2017), which was a gift from Marin Barišić and Helder Maiato (Institute for Molecular Cell Biology, University of Porto, Portugal); U2OS cell line stably expressing 2xGFP-EB3 and mCherry-CENP-A, a gift from Julie Welburn (University of Edinburgh, United Kingdom) (Kajtez et al., 2016); HeLa-Kyoto BAC lines stably expressing MKLP1-GFP, Kif4A-GFP, Kif18A-GFP, CENP-E-GFP and PRC1-GFP were a courtesy of Ina Poser and Tony Hyman (MPI-CBG, Dresden, Germany); unlabeled HeLa-TDS cells from the High-Throughput Technology Development Studio (MPI-CBG, Dresden); HeLa-TDS cells, permanently transfected with pEGFP-α-tubulin, used in our previous work (Kajtez et al., 2016); HeLa cells stably expressing YFP-tubulin, a courtesy of Lars Jansen (University of Oxford, United Kingdom); HeLa cells permanently transfected with EGFP-CLASP1, which was a gift from Helder Maiato (Institute for Molecular Cell Biology, University of Porto, Portugal). Cells were grown in flasks in Dulbecco’s Modified Eagle’s Medium (DMEM; Lonza, Basel, Switzerland) with 1 g/L D-glucose, L-glutamine, and pyruvate, supplemented with 10% of heat-inactivated Fetal Bovine Serum (FBS; Sigma Aldrich, St. Louis, MO, USA), 100 IU/mL penicillin and 100 mg/mL streptomycin solution (Lonza). For cell lines with stable expression of fluorescently labeled proteins, 50 µg/mL geneticin (Life Technologies, Waltham, MA, USA) was added. Cells were kept at 37°C and 5% CO_2_ in a Galaxy 170 R humidified incubator (Eppendorf, Hamburg, Germany). All used cell lines were confirmed to be mycoplasma free by using MycoAlert Mycoplasma Detection Kit (Lonza).

### Plasmids

To make PRC1-tgRFPt-SspB WT (opto-PRC1) plasmid, PRC1 fragment was amplified from His6-PRC1 plasmid (Addgene number: 69111) (Nixon et al., 2015) using the primers GCTAGAATTGACCGGATGAGGAGAAGTGAGGTGCTG (FWD) and CATGGTGGCGACCGGTAAATTCGAAGCTTGAGCTCGAGATCTGAGGGACTGGATGTTGGTTGAATTGAGG (REV) and inserted into plasmid tgRFPt-SspB WT (Addgene number: 60415) (Guntas et al., 2015) using *AgeI* restriction site. This step was performed using commercially available *In-Fusion HD Cloning Kit* (Clontech, Mountain View, CA, USA). The produced plasmid expresses PRC1 tagged with tgRFPt and SspB at the C-terminus. Plasmid iLID-CAAX was purchased (Addgene number: 85680) (O’Neill et al., 2016). Plasmids pEGFP-C1Kif4a-sires and EGFP-Kif18A were a gift from Jason Stumpff (University of Vermont, Burlington, VT, USA) (Stumpff et al., 2008; Stumpff et al., 2012). Plasmid GFP-CENP-E was a gift from Marin Barišić (Danish Cancer Society Research Center, Copenhagen, Denmark).

### Sample preparation

For depletion of endogenous PRC1 before opto experiments, cells were transfected 72 h (U2OS cells) or 24 h (HeLa cells) prior to imaging with 25 µL of 20 µM *Accell PRC1 siRNA* (A-019491-15-0020, Dharmacon, Lafayette, CO, USA) targeting 3’ UTR of PRC1 mRNA. A day prior to imaging, siRNA treated cells were transfected with corresponding plasmids in following amounts: 0.3 µg of iLID-CAAX, 5.5 µg PRC1-tgRFPt-SspB-WT (resistant to the used RNAi), 0.5 µg pEGFP-C1Kif4a-sires, 1 µg EGFP-Kif18A and 1 µg GFP-CENP-E. In HeLa BAC lines, 24 h prior to imaging, mock experiment cells were transfected with 100 nM *Accell Non-targeting Pool* (D-001910-10-05; Dharmacon), PRC1 siRNA treated with 100 nM *Accell PRC1 siRNA* (Dharmacon), Kif4A siRNA treated with 100 nM *Kif4A siRNA* (sc-60888; Santa Cruz Biotechnologies, Dallas, TX, USA), whereas Kif18A siRNA treated with 100 nM *Silencer Select Validated Kif18A siRNA* (s37882; Ambion, Austin, TX, USA). All transfections were performed using Nucleofector Kit R with the Nucleofector 2b Device (Lonza) using X-001 program for U2OS and O-005 (high viability) program for HeLa cells. Following transfection, the cells were seeded on 35-mm glass coverslip uncoated dishes with 0.17-mm (1.5 coverglass) glass thickness (MatTek Corporation, Ashland, MA, USA) in 1.5 mL DMEM medium with appropriate supplements. To visualize microtubules, cells were stained with silicon rhodamine (SiR)-tubulin (Lukinavicius et al., 2014) (Spirochrome AG, Stein am Rhein, Switzerland), a far-red tubulin dye, at a concentration of 50 nM 12-16 h prior to imaging. To prevent dye efflux, verapamil, a broad-spectrum efflux-pump inhibitor (Spirochrome Ltd.), was added in U2OS cells at a concentration of 1 µM. To visualize chromosomes and determine the phase of the mitosis, 20 min prior to imaging SiR-DNA (Lukinavicius et al., 2015) (Spirochrome AG, Stein am Rhein, Switzerland) was added to a final concentration of 150 nM. For experiments on U2OS cells expressing 2xGFP-EB3 and mCherry-CENP-A, the cells were synchronized by adding 20 µM of the proteasome inhibitor MG-132 (Sigma-Aldrich) to arrest the cells in metaphase. Imaging was started 15 min after adding MG-132.

### Immunocytochemistry

Cells were fixed with ice-cold methanol for 3 min, washed 3 times with phosphate buffer saline (PBS), followed by 15 min permeabilization with 0.5% Triton in PBS. Cells were washed 3 times with PBS and blocked in 1% Normal Goat Serum (NGS) exept in experiment for intensity of opto-PRC1 where cells were blocked in BSA in PBS for 1 h at 4°C. Cells were washed 3 times with PBS and then incubated in primary antibody solution in blocking buffer over night at 4°C. Following primary antibodies were used: mouse anti-PRC1 monoclonal antibody (1:100; C-1; sc-376983, Santa Cruz Biotechnology), rabbit anti-α-tubulin polyclonal antibody (1:100; SAB4500087, Sigma-Aldrich Corporation, St. Louis, MO, USA), mouse monoclonal anti-Kif4A antibody (1:100; E-8; sc-365144, Santa Cruz Biotechnology), rabbit polyclonal anti-MKLP1 antibody (1:100; N-19; sc-867, Santa Cruz Biotechnology), mouse monoclonal anti-Eg5 antibody (1:100; A-1; sc-365681, Santa Cruz Biotechnology). After washing of primary antibodies with PBS, cells were incubated in a solution of secondary antibodies in 2% NGS or BSA in PBS for 1 h at room temperature protected from light. Following secondary antibodies were used: donkey anti-mouse IgG Alexa Fluor 488 (1:250; ab150109, Abcam, Cambridge, UK), donkey anti-rabbit IgG Alexa Fluor 594 (1:250; ab150064, Abcam), donkey anti-rabbit IgG Alexa Fluor 405 (1:250; ab175649, Abcam), and goat anti-mouse IgG Alexa Fluor 647 (1:250; ab150119, Abcam). After washing of the secondary antibodies 3 times in PBS, cells were incubated with a solution of 4’,6-diamidino-2-phenylindole (DAPI) (1:1000) in PBS for 20 min and washed 3 times in PBS or SiR-DNA (150 nM) in PBS for 15 min before imaging. Note that we used immunocytochemistry for PRC1 rather than Western blot analysis because the efficiency of opto-PRC1 plasmid transfection is low, and as Western blot analysis provides information about the complete cell population, these results may not be relevant for the cells used in the optogenetic experiments. In contrast, by using immunocytochemistry we analyzed only the cells with a similar opto-PRC1 level and in the same phase as those in our optogenetic experiments.

### Microscopy

Immunocytochemistry imaging and live imaging of unlabeled U2OS, U2OS stably expressing CENPA-GFP, HeLa-TDS pEGFP-α-tubulin and HeLa BAC CENP-E-GFP cells was performed using Bruker Opterra Multipoint Scanning Confocal Microscope (Bruker Nano Surfaces, Middleton, WI, USA), described previously (Buda et al., 2017). In brief, the system was mounted on a Nikon Ti-E inverted microscope equipped with a Nikon CFI Plan Apo VC 100x/1.4 numerical aperture oil objective (Nikon, Tokyo, Japan). During imaging, live cells were maintained at 37°C using H301-K-frame heating chamber (Okolab, Pozzuoli, NA, Italy). In order to obtain the optimal balance between spatial resolution and signal-to-noise ratio, 60-µm pinhole aperture was used. Opterra Dichroic and Barrier Filter Set 405/488/561/640 was used to separate the excitation light from the emitted fluorescence. Following emission filters were used: BL HC 525/30, BL HC 600/37 and BL HC 673/11 (all from Semrock, Rochester, NY, USA). Images were captured with an Evolve 512 Delta Electron Multiplying Charge Coupled Device (EMCCD) Camera (Photometrics, Tucson, AZ, USA) using a 200 ms exposure time. Electron multiplying gain was set on 400. Camera readout mode was 20 MHz. No binning was performed. The xy-pixel size in the image was 83 nm. The system was controlled with the Prairie View Imaging Software (Bruker).

For kinetics experiments on U2OS cells (**Fig. 1**), 561 and 488 nm diode laser lines were used every 10 s with 200 ms exposure time. In all other optogenetic experiments, stacks were acquired using sequentially the following diode laser lines: 561 nm (to visualize opto-PRC1), 488 nm (to activate the optogenetic system and to visualize GFP), and 640 nm (to visualize SiR-tubulin or SiR-DNA, when applicable), with time interval between z-stacks of 60 s and with 200 ms exposure time per laser line. To prevent dissociation of PRC1 from the cell membrane between acquiring two consecutive z-stacks, only blue light was turned on for 200 ms every 10 seconds. Cells were imaged this way for 20 min in order to achieve almost complete removal of PRC1 from the spindle, after which the blue light was turned off and imaging was continued for another 10 min at 60 s intervals. The total imaging time of 30 min was chosen to be close to the typical metaphase duration of 29.7 ± 2.3 min, which was measured from the metaphase plate formation until anaphase onset in U2OS cells expressing CENP-A-GFP, mCherry-α-tubulin and PA-GFP-tubulin (N=187) imaged after nuclear envelope breakdown every minute by obtaining 15 z-slices with 0.5 µm spacing and 150 ms exposure time. After 30 min of imaging, one z-stack in each of the three channels was taken in order to visualize the spindle and kinetochores after PRC1 return. In all cells except HeLa cells expressing pEGFP-α-tubulin, three focal planes with spacing between adjacent planes of 1 µm were acquired. Live imaging of HeLa cells stably expressing pEGFP-α-tubulin was performed in the same manner as described above for optogenetic experiments. Additionally, before turning the blue light on every 10 s, one z-stack was acquired using 561, 488 and 640 nm diode laser lines with averaging 8, and seven focal planes with spacing between adjacent planes of 0.5 µm. Stack was taken in the same way after 20 min of exposure to the blue light and again 10 min after the blue light was switched off. Imaging of HeLa BAC CENP-E-GFP mock and PRC1 siRNA treated cells was performed by acquiring one z-stack of 3 focal planes with spacing between adjacent planes of 1 µm. For imaging of immunostained cells, 5 focal planes with spacing between adjacent planes of 0.5 µm were acquired.

Live imaging of U2OS cells stably expressing 2x-GFP-EB3 and mCherry-CENP-A was performed on a spinning disk confocal microscope system (Dragonfly, Andor Technology, Belfast, UK) using 63x/1.47 HC PL APO oil objective (Leica) and Zyla 4.2P scientific complementary metal oxide semiconductor (sCMOS) camera (Andor Technologies). During imaging cells were maintained at 37° and 5% CO2 within H301-T heating chamber (Okolab, Pozzuoli, Italy). Images were acquired using Fusion software (v 2.2.0.38). For excitation, 488-nm and 561-nm laser lines were used for visualisation of GFP, and mCherry and opto-PRC1, respectively. In order to achieve PRC1 removal from the spindle, 3 z-planes with a z-spacing of 1 µm were acquired sequentially with both laser lines, every 10 s with 200 ms exposure time for 10 min. This imaging protocol was followed by faster imaging, every 1.5 s with both laser lines on a central z-plane in order to visualize EB3 dynamics. Control, mock and PRC1 siRNA treated cells were imaged with the same imaging protocol.

Live imaging of unlabeled, BAC (except CENP-E-GFP), YFP-tubulin and EGFP-CLASP1 HeLa cell lines was performed on Leica TCS SP8 X laser scanning confocal microscope with a HC PL APO 63x/1.4 oil immersion objective (Leica, Wetzlar, Germany) heated with an objective integrated heater system (Okolab, Burlingame, CA, USA). During imaging, cells were maintained at 37°C in Okolab stage top heating chamber (Okolab, Burlingame, CA, USA). The system was controlled with the Leica Application Suite X software (LASX, 1.8.1.13759, Leica, Wetzlar, Germany). For GFP- and YFP-labeled proteins, a 488-nm and 514-nm Argon laser was used, respectively, and for SiR-DNA or SiR-tubulin, a 652-nm white light laser was used. For AF-647, a 637-nm white light laser was used. GFP, and SiR-DNA, SiR-tubulin or AF-647 emissions were detected with hybrid detector. For mock, PRC1 siRNA, Kif4A siRNA, and Kif18A siRNA experiments, images were acquired at 1–3 focal planes with 1 µm spacing and 0.05-µm pixel size. In optogenetic experiments 3 z-stacks with 1 µm spacing were acquired sequentially every 10 s in the same manner as in optogenetic experiments in U2OS cells. One z-stack with line averaging of 6 or 16 was acquired before system activation, 20 min after exposure to blue light and 10 min after the light was switched off.

### Image and data analysis

Since the cells were transiently transfected with opto-PRC1, we observed variability in PRC1 expression levels and therefore we imaged and analyzed only those metaphase spindles with PRC1 localization consistent with endogenous and fluorescently labeled PRC1 (Kajtez et al., 2016; Polak et al., 2017). Cells were not synchronized in order to avoid additional chemical treatment of cells, and metaphase was determined by alignment of kinetochores in the equatorial plane.

For determination of kinetics of PRC1 removal and return (**Fig. 1 C**), intensity of opto-PRC1 was measured in each time frame on one focal plane. We used *Poligon selection* tool in Fiji (National Institutes of Health, Bethesda, MD, USA) to encompass the area of the spindle, *A_spindle_*, and measure mean spindle intensity, *M_spindle_*. Mean background intensity in the cytoplasm was measured using 30 x 30 pixels rectangle, *M_cyto_*. Spindle intensity was background corrected by subtracting *M_cyto_* from *M_spindle_* to obtain *M_spindle corr_*. In order to calculate the sum of PRC1 intensity on the spindle, *M_spindle corr_* was multiplied with *A_spindle_* for each timeframe. The background intensity outside of the cell was negligible, thus we did not take it into account. Note that for the measurements of kinetic parameters in **Fig. 1 C**, four outliers were excluded (see **Fig. S1 E**). The percentage of PRC1 removal was calculated from the mean value of intensity of all cells at time 20 min, i.e., the last time point of the continuous exposure to the blue light. The percentage of return was calculated from the mean value of intensity of all cells in the interval 25-30 min, i.e., during last 5 min of PRC1 return.

Intensity profiles of opto-PRC1 removal and return (**Fig. 1 D**) were obtained on sum intensity projections of all three z-planes by drawing a pole-to-pole line along the long axis of the spindle by using *Line* tool in Fiji. The width of the line corresponded to the width of each individual spindle. Intensities were normalized to position of the poles.

For quantification of PRC1 knock-down by siRNA and intensity level of opto-PRC1 (in **Fig. S1 A,B**) PRC1 intensity on fixed cells was measured on a sum-intensity projection of five focal planes by the procedure described above, in a way that mean spindle intensity was background corrected by subtracting mean intensity in the cytoplasm. For measuring opto- PRC1 intensity on the spindle cells where PRC1 was visible on astral microtubules were not analyzed, nor imaged in live experiments.

The GFP-tubulin signal intensity of a cross-section of a bridging fiber was measured by drawing a 3-pixel-thick line between and perpendicular to the tubulin signal joining the tips of sister k-fibers. The intensity profile was taken along this line and the mean value of the background signal present in the cytoplasm was subtracted from it. The signal intensity of the bridging fiber was calculated as the area under the peak using SciDavis (Free Software Foundation Inc., Boston, MA, USA). The signal intensity in the region lateral from k-fiber tip was measured in a similar manner, 1 µm away from a k-fiber tip, perpendicular to and crossing the corresponding k-fiber. All of the measurements were performed on a single z-plane. The profile intensity of bridging fiber and at the position lateral from k-fiber tip in SiR-tubulin (**Fig. 3 B,C**) was performed in the same manner. Note that mostly outermost fibers were used for these measurements because of being most easily distinguished from neighboring fibers.

Shapes of the spindle were quantified by tracking outermost k-fiber contours in the central z-slice of the spindle, or maximum intensity z-projection of two central z-slices. All spindles were rotated to have horizontal long axis. Pole-to-k-fiber-tip tracking was done using *Multipoint* tool by placing 5 roughly equidistant points along contour length, first point being at the pole and the last point being at the k-fiber tip. First point of each contour was translated to (0,0). This was done for maximum of 4 trackable outermost k-fibers in the spindle. Curvature of the contour was calculated by fitting a circle to the contour points of individual k-fibers and retrieving reciprocal value of its radius. To test the effect of tracking errors on curvature, we introduced 1-pixel noise to the x and y values of tracked points, which did not change the result. Angle between outermost k-fibers (θ) was calculated as the angle between lines passing through last two points along the contour of sister k-fibers.

Spindle length and width were measured using *Line* tool in Fiji. For length measurements, a line was drawn from pole to pole. The position of the pole was defined as the location of the strongest tubulin signal. Width was measured by drawing a line between outermost kinetochore pairs on the opposite sides of the spindle, perpendicular to the spindle long axis.

EB3 spots in U2OS cells stably expressing 2x-GFP-EB3 and mCherry-CENP-A were tracked by obtaining their xy coordinates using *Multipoint* tool in Fiji from the frame when a spot appeared until it disappeared or was no longer clearly distinguishable from its neighbors. Only spots belonging to bridging fibers were traced, which were defined as those passing between sister kinetochores or moving along PRC1 streaks. Half-overlap length was measured as the distance between the last location where a tracked EB3 spot was visible and the spindle equator. The number of spots in time was obtained by visually inspecting time frames where individual kinetochore pair or PRC1 bundle was visible, and dividing the total number of the observed EB3 spots in the bridging fibers by the total time of observation of individual kinetochore pairs or PRC1 bundles. Kymographs were generated using KymographBuilder plug-in in Fiji, and half-overlap length was measured from kymographs as the distance between a spot’s trajectory end point and the mid-line between the poles corresponding to the equatorial plane. These measurements were performed in the first 2 min of a fast imaging sequence, which followed after the 10 min imaging protocol required for PRC1 removal.

Inter-kinetochore distance was measured using *Line* tool in Fiji on individual or maximum intensity z-projections of up to two z-planes as a distance between centers of sister kinetochore signals. Peripheral kinetochores were defined as three outermost pairs on each side of the spindle with respect to spindle long axis, while the remaining were considered as central. Measurement of inter-kinetochore distances in prometaphase was performed on U2OS cells expressing CENP-A-GFP, mCherry-α-tubulin and PA-GFP-tubulin, just after nuclear envelope breakdown in one imaging plane where both sister kinetochores could be clearly distinguished. For measuring kinetochore alignment and orientation, as well as orientation and length of PRC1 bundles, *Multipoint* tool in Fiji was used. A point was placed in the center of signal of each sister kinetochore or edges of PRC1 signal for each bundle. Before measuring, images were rotated in order to achieve perpendicular direction of the equatorial plane with respect to x-axis. The equatorial plane was defined with two points placed between outermost pairs of kinetochores on the opposite sides of the spindle. For all measurements regarding kinetochores, those located at the spindle poles were not taken into account. All measurements in opto and control cells were performed in three time-points: before the blue light was switched on, after 20 min of continuous exposure to the blue light, and 10 min after the blue light was turned off. In untreated and PRC1 siRNA treated cells measurements were performed at the beginning of imaging. Kymographs of kinetochore oscillations were produced by Low Light Tracking Tool (LLTT), an ImageJ plugin (Krull et al., 2014). Tracking of kinetochores in x, y plane was performed on maximum intensity of 2-3 z-planes. Sigma value (standard deviation of the Gaussian used to approximate the Point Spread Function (PSF) of the tracked objects) was set to *Automatic*.

The intensity of GFP-Kif4A in metaphase and anaphase was measured in GFP-Kif4A channel on sum-intensity projections of all three z-planes using *Poligon selection* tool in Fiji by encompassing chromosomes in SiR-DNA channel. The intensity of opto-PRC1 in metaphase was measured by encompassing the spindle area. Background cytoplasm intensities in both GFP-Kif4A and opto-PRC1 channels were measured in 30 x 30 pixels rectangle and subtracted from measured mean intensities of GFP-Kif4A and opto-PRC1. Corrected intensities were divided by the number of focal planes. Measurements were performed in three time-points in metaphase: before the blue light was switched on, after 20 min of continuous exposure to the blue light, and 10 min after the blue light was turned off. In anaphase, measurements were performed 4 min after anaphase onset. Anaphase onset was defined as the timeframe when separation of majority of sister chromatids in SiR-DNA channel occurred.

The mean signal intensities of opto-PRC1 and MKLP1-GFP were measured on the central z-plane using *Poligon selection* tool in Fiji by encompassing the spindle area in SiR-tubulin channel. Background cytoplasm intensities in both channels were measured in 30 x 30 pixels rectangle and subtracted from measured mean intensities of opto-PRC1and MKLP1-GFP. Measurements were performed in the same time-points as for Kif4A in metaphase.

For quantification of PRC1 knock-down by siRNA in HeLa BAC MKLP1-GFP cell line, PRC1 intensity was quantified in the same manner as in U2OS cells. The intensity levels of MKLP1-GFP and Kif4A-GFP in fixed HeLa cells were measured by encompassing the whole cell area using *Poligon selection* tool in Fiji on sum intensity projections of 5 z-planes. Localization test of GFP-Kif4A or Kif4A-GFP, MKLP1-GFP, Kif18A-GFP, EGFP-CLASP1, and CENP-E-GFP in the bridging fibers of either opto cells or cells treated with mock siRNA or PRC1 siRNA was performed by visually inspecting the GFP signal through the z-stack, in the region where PRC1-labeled fibers were found, i.e., in the region that spans between sister kinetochores and continues ∼2 µm laterally from sister kinetochores. Quantification of the signal length of PRC1-GFP in mock, Kif4A, and Kif18A siRNA treated HeLa cells was performed by drawing a 5-pixel-thick pole-to-pole line along individual bundles and calculated as the width of the peak of the PRC1-GFP signal intensity.

Statistical analysis was performed in MATLAB (MathWorks, Natick, MA, USA) and RStudio (R Foundation for Statistical Computing, Vienna, Austria). Normality of the data was inspected by Q-Q plots and Shapiro-Wilk test of normality. Normally distributed data were tested with two-tailed t-test, while Mann-Whitney test was used for non-normally distributed data, as noted in figure captions. Proportions were statistically compared with test for equality of proportions, two-proportions z-test. For data with expected count smaller than 5, Yates’s correction for continuity was used.

Graphs were generated in MATLAB (MathWorks) and RStudio (R Foundation for Statistical Computing). ImageJ (National Institutes of Health, Bethesda, MD, USA) was used to crop and rotate images, and to adjust brightness and contrast to the entire image, which was applied equally to all images in the same panel. To remove high frequency noise in displayed images a Gaussian blur filter with a 0.5-pixel sigma (radius) was applied where stated. Figures were assembled in Adobe Illustrator CS5 (Adobe Systems, Mountain View, CA, USA).

## Acknowledgements

We thank Helder Maiato and Marin Barišić for unlabeled U2OS and U2OS CENP-A-GFP cell lines; Julie Welburn for U2OS 2xGFP-EB3 mCherry-CENP-A cell line; Ina Poser and Tony Hyman for HeLa MKLP1-GFP, Kif4A-GFP, Kif18A-GFP, CENP-E-GFP and PRC1-GFP cell lines; Helder Maiato for HeLa EGFP-CLASP1 cell line; Lars Jansen for HeLa YFP-tubulin cell line; Marin Barišić for GFP-CENP-E plasmid; Jason Stumpff for pEGFP- C1Kif4a-sires and EGFP-Kif18A plasmids; Nenad Pavin, Agneza Bosilj, Kruno Vukušić, and Juraj Simunić for discussions and constructive comments on the manuscript; Sonja Lesjak for help with the cloning and plasmids; Ivana Šarić for the drawings. This work was funded by European Research Council (ERC, GA number 647077) and Croatian Science Foundation (HRZZ, project IP-2014-09-4753).

## Author Contributions

M.J, P.R. and J.M. performed and analyzed all experiments, and wrote the manuscript together with I.M.T. A.M. made the optogenetic plasmid and performed pilot experiments. I.M.T. conceived and supervised the project.

## Data availability

All data are available from the corresponding author upon request.

## Code availability

Custom codes used in this study are available upon request.

## Supplementary Figures

**Figure S1.**
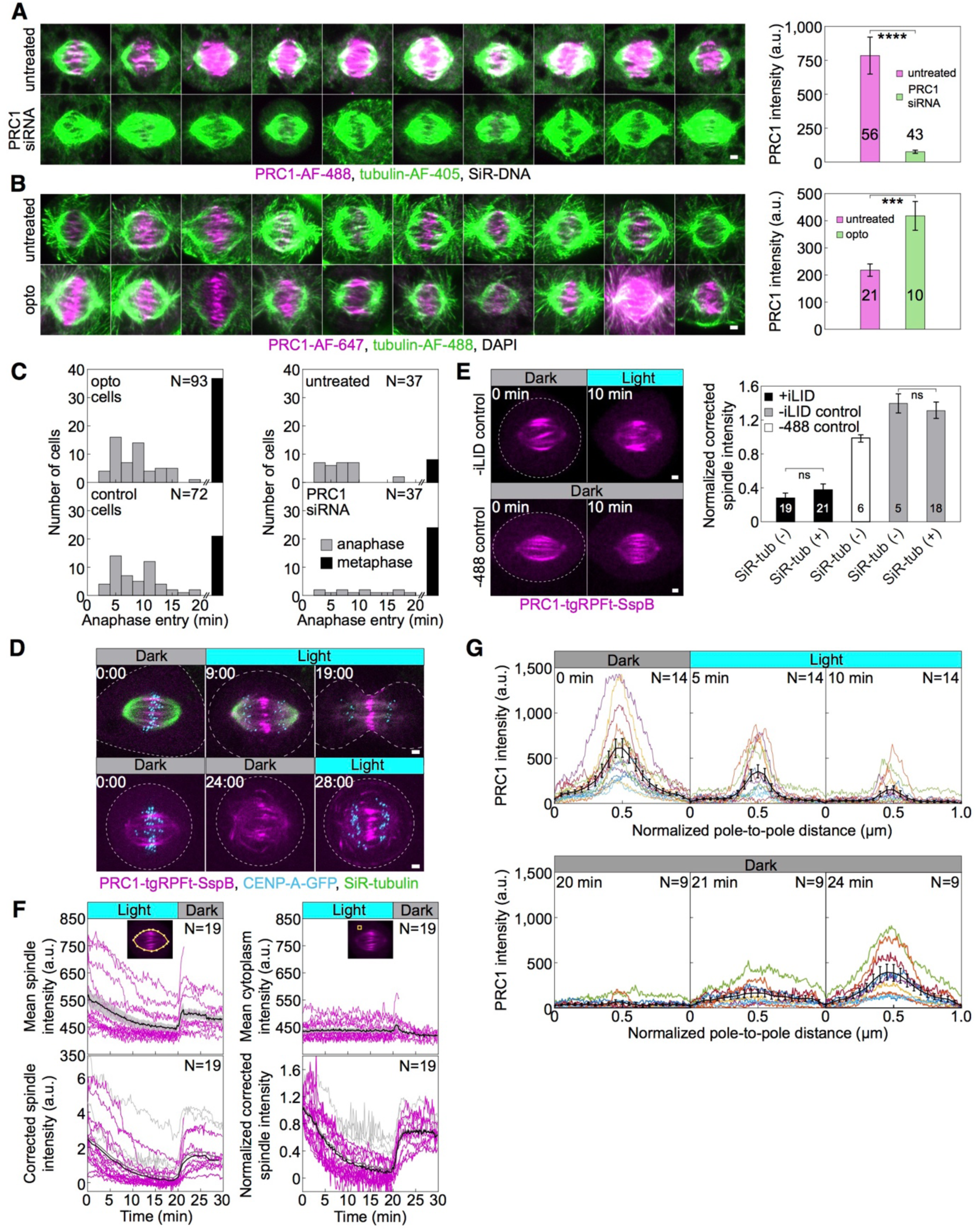
(**A**) Metaphase spindles in fixed, unlabeled U2OS cells (left) immunostained for endogenous PRC1 (AF-488, magenta), tubulin (AF-405, green) and stained with SiR-DNA (not shown) in untreated (top) and PRC1 siRNA treated cells (bottom). All images are sum intensity projections of five z-planes. Graph (right) shows PRC1 intensity in untreated (magenta) and PRC1 (green) siRNA treated cells. Data obtained from three independent experiments. (**B**) Metaphase spindles in fixed, unlabeled U2OS cells immunostained for PRC1 (AF-647, magenta), tubulin (AF-488, green) and stained with DAPI (not shown) in untreated (top) and PRC1 siRNA treated cells with addition of opto-PRC1 (bottom). All images are sum intensity projections of five z-planes. Graph (right) shows PRC1 intensity in untreated (magenta) and cells with depleted endogenous and added opto-PRC1 (green). Data obtained from two independent experiments. (**C**) Number of cells that entered anaphase (gray bars) and those remained in metaphase (black bars) during 20 min of exposure to the blue light for opto and control cells (left), and untreated and PRC1 siRNA treated cells (right). For opto and control cells those imaged for measurements of SiR-DNA area are also taken into account. (**D**) Time-lapse images of U2OS cells with stable expression of CENP-A-GFP (cyan) where control cell (top) progresses to anaphase during and opto cell (bottom) after 20-min exposure to the blue light. (**E**) Time-lapse images of U2OS cells with stable expression of CENP-A-GFP (not shown) and transient expression of opto-PRC1 and iLID-CAAX, non-exposed to the blue light (-488 control, bottom left) and exposed to blue light but without iLID-CAAX (-iLID control, top left), before and after 10 minutes of imaging. Graph (right) shows opto-PRC1 intensities after 10 min of imaging in +iLID cells with and without SiR-tubulin (black), -488 control cells (white) and –iLID control cells with and without SiR-tubulin (gray). Note that SiR-tubulin does not affect opto-PRC1 removal. (**F**) Raw opto-PRC1 intensities from individual cells (magenta lines) at the metaphase spindle (top left) and in cytoplasm (top right) during removal and return to the spindle. Mean (thick black line) and s.e.m. (shaded area). Opto-PRC1 intensities at the metaphase spindle corrected for cytoplasm (bottom left) and normalized to the initial value (bottom right). Outliers (gray lines), mean without outliers (thick black line), mean with outliers (thick gray line). Note that outliers are excluded in **Fig. 1 C**. (**G**) Pole-to-pole opto-PRC1 intensity during removal (top) and return (bottom) to the spindle as in **Fig. 1 D**. Individual cells (colored lines), mean (thick black line), s.e.m (error bars). Images are max projection of 3 z-planes. N and numbers inside bars; number of cells. Scale bars; 2 µm. Statistical analysis; t-test. p-value legend as in **Fig. 2**.

**Figure S2.**
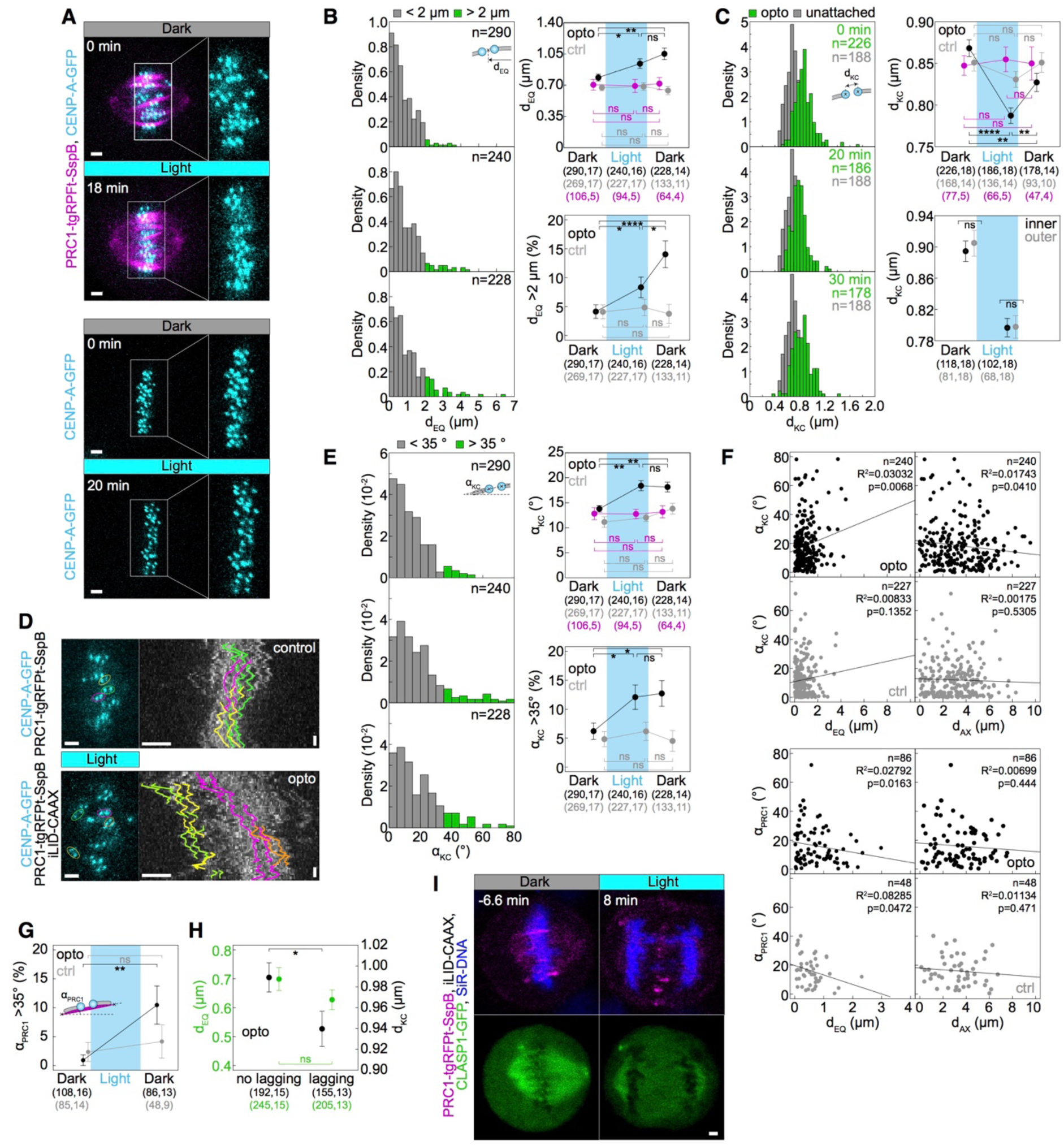
(**A**) Spindle in a U2OS cell stably expressing CENP-A-GFP (cyan) with (top) and without (bottom) transient expression of opto-PRC1 (magenta) before (0 min, Dark) and at the end of continuous exposure to the blue light (18 min, bottom). Enlargements of the boxed regions show kinetochores only. Images are maximum intensity projections of three z-planes, smoothed with 0.5-pixel Gaussian blur. (**B**) Density histograms (left) of the distance from the midpoint between sister kinetochores to the equatorial plane, d_EQ_, for opto U2OS cells before (0 min, top), at the end of continuous exposure (20 min, middle) and 10 min after cessation of exposure to the blue light (30 min, bottom). d_EQ_ greater (green) and lesser than 2 µm (gray) is shown. This corresponds to roughly 95^th^ percentile of opto data before system activation. Percentage of d_EQ_ greater than 2 µm in opto (black) and control (gray) cells (bottom right). Graph (top right) showing d_EQ_ in control (gray) and opto cells with and without SiR-tubulin (black), and cells only expressing CENP-A-GFP (magenta) in corresponding timepoints. (**C**) Density histograms (left) of inter-kinetochore distance, d_KC_, for opto U2OS cells (green) for time-points as in **B**, overlaid on density histogram of unattached d_KC_ from early prometaphase cells (mean: 0.66 ± 0.01 µm, N=7, n=188) (gray). Inter-kinetochore distance (d_KC_) (top right) in control (gray) and opto cells with and without SiR-tubulin (black), and cells only expressing CENP-A-GFP (magenta) in same timepoints. Inter-kinetochore distance (d_KC_) (bottom right) of inner (black) and outer (gray) kinetochore pairs before and 20 min after exposure to the blue light. (**D**) Metaphase plates (left) of control (top) and opto (bottom) U2OS cell expressing CENP-A-GFP (cyan). Corresponding kymographs (right) show kinetochore fluctuations during last 10 min of 20-min exposure to the blue light, when there was no PRC1 on the spindle in opto cells. Kinetochore tracks (right) are color-coded with respect to corresponding sister pairs (left). Note that in opto cell (bottom) the displaced kinetochores (yellow, green) showed similar extent of oscillations as non-displaced kinetochores in opto (bottom) and all in control (top) cells (N=10 cells in each condition). Vertical scale bar; 1 min. (**E**) Density histogram (left) of angle between sister kinetochore axis and the long spindle axis, α_KC_, for opto U2OS cells for time-points as in **B**. α_KC_ greater (green) and lesser than 35° (gray) is shown. This corresponds to roughly 95^th^ percentile of opto data before system activation. Percentage of α_KC_ greater than 35° in opto (black) and control (gray) cells (bottom right). Graph (top right) showing α_KC_ in control (gray) and opto cells with and without SiR-tubulin (black), and cells only expressing CENP-A-GFP (magenta) in timepoints as in **B**. (**F**) Graphs show α_KC_ versus corresponding d_EQ_ and the distance from the midpoint between sister kinetochores to the long spindle axis, d_AX_ (left), and α_PRC1_ versus d_EQ_ and d_AX_ (right) in opto (top, black) and control (bottom, gray) cells. Kinetochore parameters correspond to 20 min after continuous exposure to the blue light, and PRC1 parameters correspond to 10 min after cessation of exposure. Note that PRC1 is not on the spindle at 20 min in opto cells. Black lines show linear regression. (**G**) Percentage of angles between the line connecting the end points of the PRC1 streak and the long spindle axis, α_PRC1_, greater than 35° in opto (black) and control (gray) cells before exposure and 10 min after cessation of exposure to the blue light. (**H**) d_KC_ (black) and d_EQ_ (green) in opto anaphase cells with and without lagging kinetochores. (**I**) Time-lapse images of HeLa cell stably expressing EGFP-CLASP1 with transient expression of opto-PRC1 (magenta) and iLID-CAAX, and stained with SiR-DNA (blue). Top (merge tgRFPt and SiR-DNA), bottom (GFP). Images are maximum intensity projections of 3 z-planes, smoothed with 0.5-pixel Gaussian blur. Anaphase onset is set as time 0 min. Note the absence of GFP signal in the spindle midzone and lagging chromosomes at 8 min. Cyan rectangles inside graphs indicate exposure to the blue light. Numbers in brackets; number of measurements and cells, respectively. Time; min. n; number of measurements. Horizontal scale bars; 2 µm. Statistical analysis; two-proportions z-test. R^2^, coefficient of determination. p-value legend as in **Fig. 2**.

**Figure S3.**
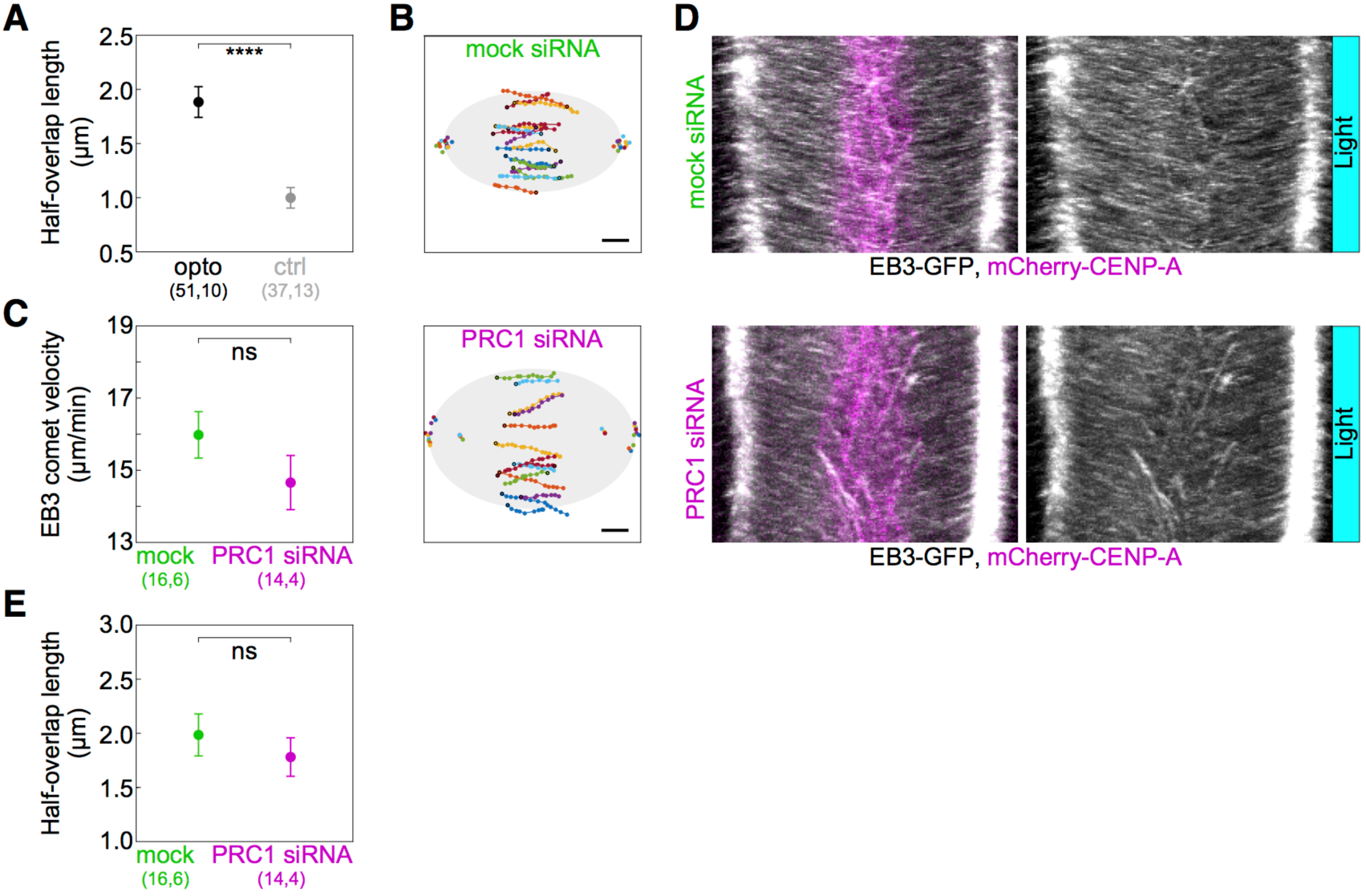
**(A)** Half-overlap length in opto (black) and control (gray) U2OS cells expressing 2xGFP-EB3 measured from kymographs as in **Fig. 3 E**, as the distance of spot’s track end-point to the equatorial plane. All measurements were performed within 2 min following the 10 min imaging protocol required for removal of PRC1. (**B)** Trajectories of all tracked EB3 spots (connected dots) in mock (top) and PRC1 siRNA (bottom) treated U2OS cells expressing 2xGFP-EB3. Black dot; start of trajectory. Single dots on the left and the right side; spindle poles. (**C**) Velocity of EB3 spots within the bridging fiber in mock (green) and PRC1 siRNA treated (magenta) cells. (**D**). Kymographs of mock (top) and PRC1 siRNA (bottom) treated U2OS cells expressing 2xGFP-EB3 after 10 min of imaging protocol required for removal of PRC1. Left (merge opto-PRC1, mCherry and GFP), right (GFP). (**E**) Half-overlap length in mock (green) and PRC1 siRNA (magenta) treated U2OS cells expressing 2xGFP-EB3 measured as in **Fig. 3G** and retrieved from tracks showed in **B**. Filled cyan rectangles indicate exposure to the blue light. Numbers in brackets denote measurements and cells; single numbers denote cells. Error bars; s.e.m. Scale bars; 2 µm. Statistical analysis; t-test. p-value legend as in **Fig. 2**.

**Figure S4.**
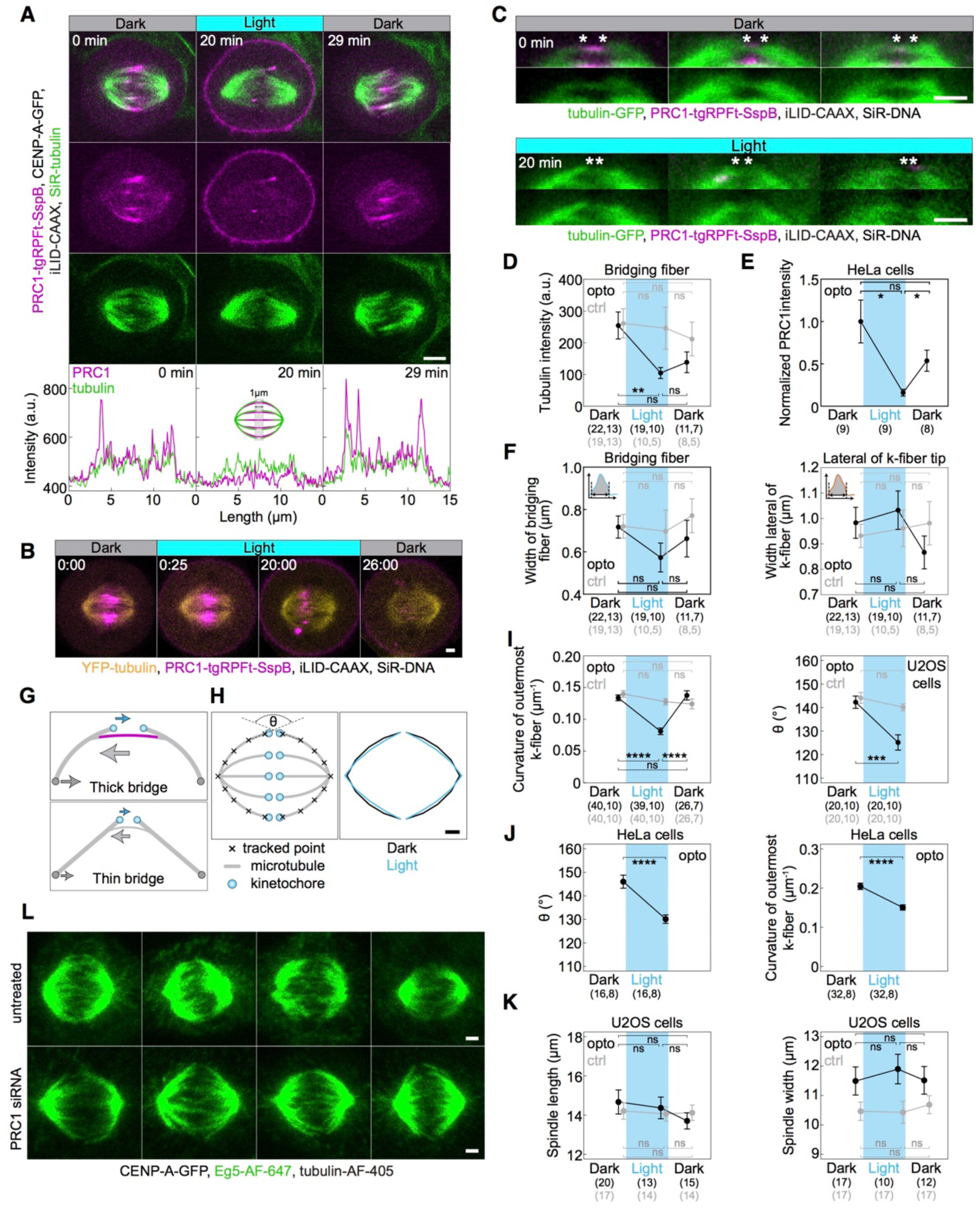
**(A)** Spindle in a U2OS cell stably expressing CENP-A-GFP (not shown), with transient expression of opto-PRC1 (magenta) and iLID-CAAX, and stained with SiR-tubulin (green) before (0 min, Dark), at the end of continuous exposure (20 min, Light) and 9 min after cessation of exposure to the blue light (29 min, Dark). Top (merge), middle (tgRFPt), bottom (SiR). Graphs (bottom) correspond to signal intensity profiles of opto-PRC1 (magenta) and SiR-tubulin (green) taken across equatorial plane in shown spindles. Profile intensities were acquired on individual z-planes using a 1-µm thick line across the spindle midzone. Images are individual z-plane, smoothed with 0.5-pixel-sigma Gaussian blur. Scale bar; 5 µm. **(B)** Time-lapse image of a HeLa cell stably expressing YFP-tubulin (yellow), transiently opto-PRC1 (magenta) and iLID-CAAX and stained with SiR-DNA dye (not shown). In 0:00 (Dark), 561 (opto-PRC1), and 514 (tubulin) + 652 (SiR-DNA) nm lasers are used stack-sequentially. From 0:25-20:00 (Light), 561 and 488+514+652 nm lasers are used stack-sequentially, every 25 s. Note that in 0:00 min, opto-PRC1 is not present on the membrane as in **Fig. 1 B**. However, in the first frame of Light sequence, 0:25, weak signal of opto-PRC1 is present on the membrane even though opto-PRC1 was imaged in stack before 488 nm laser was on. Also, opto-PRC1 is visible on the membrane 6 min after (26:00, Dark) 488 nm laser was turned off. This is a consequence of 514 nm laser, used to visualize YFP, being able to activate optogenetic system. (**C**) Enlargements of spindles in HeLa cells with stable expression of tubulin-GFP (green), depleted for endogenous PRC1, with transient expression of opto-PRC1 (magenta) and iLID-CAAX, and stained with SiR-DNA (not shown), before exposure to the blue light (0 min, Dark, top; first row: merge, second row: GFP) and at the end of continuous exposure to the blue light (20 min, Light, bottom; first row: merge, second row: GFP). Note that at 20 min opto-PRC1 is absent from the spindle. Images do not belong to the same cell. Asterisks mark the position of kinetochores. All images are single z-plane smoothed with 0.5-pixel-Gaussian blur. Scale bar; 2 µm. (**D**) Tubulin-GFP signal intensity of the bridging fibers in opto (black) and control (gray) HeLa cells before (0 min, Dark), at the end of continuous exposure (20 min, Light) and 10 min after cessation of exposure to the blue light (30 min, Dark). Signal intensity is measured as the area under the peak shown in **Fig. 4 E**. (**E**) Graph showing normalized PRC1 intensity on the spindle in opto HeLa tubulin-GFP cells used for measurements in **D**. (**F**) Width of the bridging fiber (left) and microtubule bundle lateral from k-fiber tip (right) in opto (black) and control (gray) HeLa cells at time-points as in **D**, measured as shown in schemes (upper left corners). (**G**) Top: Schematic of force-balance in the spindle where compression in the bridging fiber (green arrow) balances compression at the poles (gray arrow) and tension at kinetochores (cyan arrow). Bottom: Upon PRC1 removal, reduction of number of microtubules in the bridging fiber is expected to reduce compression in the bridging fiber and at the poles, leading to k-fiber straightening. (**H**) Outermost k-fiber contours in opto U2OS cells before (black, Dark) and at the end of continuous exposure to the blue light (cyan, Light) plotted by connecting mean positions of all tracked points. Scheme (left) depicts how contours were tracked. θ; angle between outermost k-fibers (**I**) Curvature of the outermost k-fibers (left) in opto (black) and control (gray) U2OS cells in same time-points as in B. Angle between outermost k-fibers (right) in opto (black) and control (gray) U2OS cells at first two time-points as in B. (**J**) Angle between (left) and curvature (right) of the outermost k-fibers in HeLa cells. (**K**) Spindle length (left) and width (right) in opto (black) and control (gray) U2OS cells at timepoints as in **D**. (**L**) Metaphase spindles in fixed U2OS cells stably expressing CENP-A-GFP (not shown) immunostained for Eg5 (AF-647, green) and tubulin (AF-405, not shown) in untreated (top) and PRC1 siRNA treated cells (bottom). All images are sum intensity projections of five z-planes. Scale bar; 2 µm. Cyan rectangles inside graphs indicate exposure to the blue light. Numbers in brackets; number of measurements and cells, respectively. Statistical analysis; t-test. p-value legend as in **Fig. 2**.

**Figure S5.**
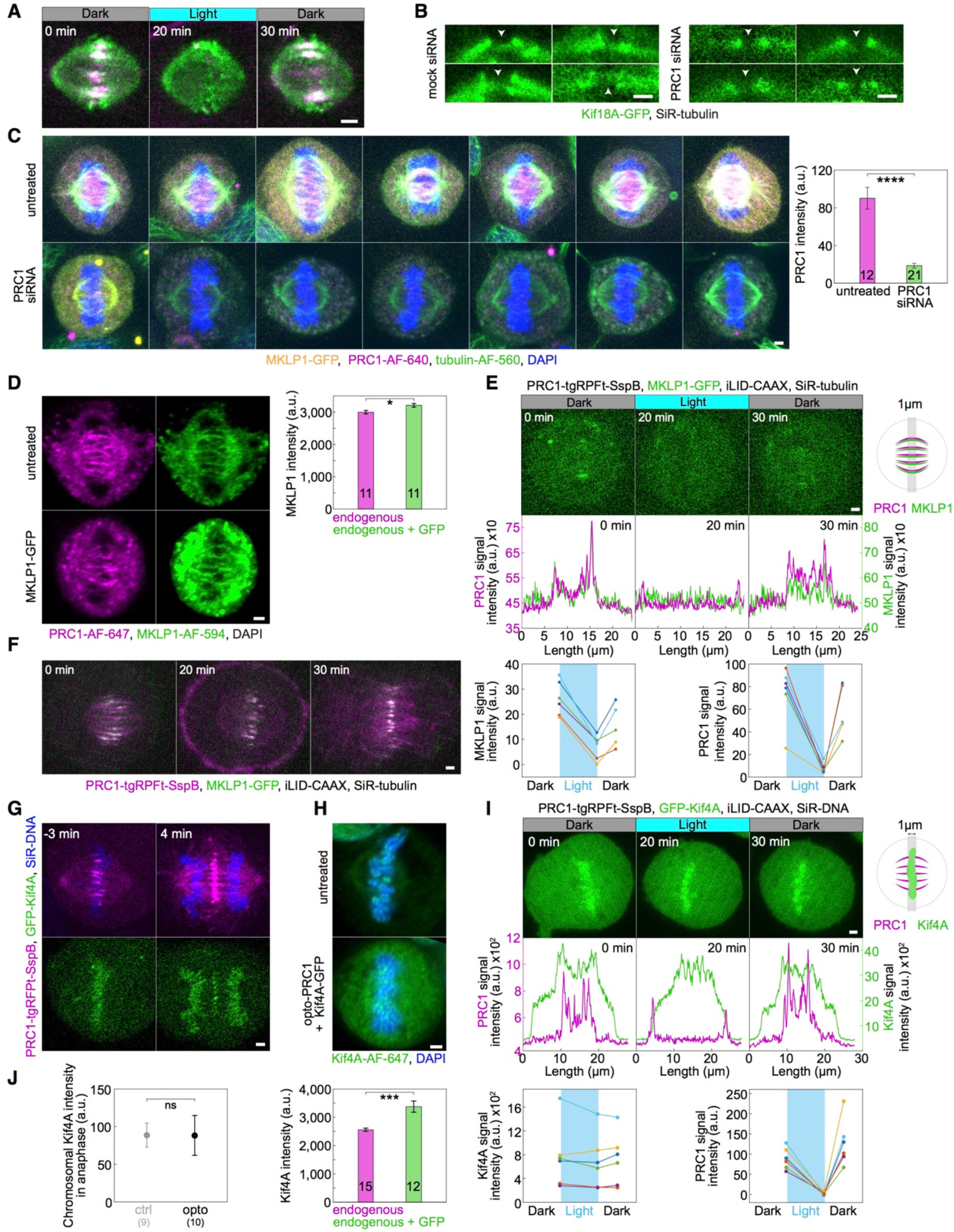
(**A**) Time-lapse of unlabeled U2OS cell with transient expression of opto-PRC1 (magenta), iLID-CAAX and EGFP-Kif18A (green) as in **Fig. 5 B**. Note the co-localization of opto-PRC1 and EGFP-Kif18A in the bridging fibers. (**B**) Enlargements of control (left) and PRC1 siRNA treated (right) HeLa BAC cells expressing Kif18A-GFP (green). Note the presence of Kif18A-GFP in the bridging fibers of control cells, and its absence in PRC1 siRNA treated cells (white arrowheads). (**C**) Spindles in fixed HeLa cells stably expressing MKLP1-GFP (yellow) and immunostained for PRC1 (AF-640, magenta), tubulin (AF-560, green) and stained with DAPI (blue) in untreated (top) and PRC1 siRNA treated (bottom) cells (left). All images are sum intensity projections of five z-planes. Graph (right) shows PRC1 intensity in untreated (magenta) and PRC1 siRNA treated (green) cells. Numbers inside bars; number of cells. (**D**) Spindle in a fixed unlabeled untreated HeLa cell (top left) and HeLa BAC cell expressing MKLP1-GFP (bottom left), immunostained for MKLP1 (AF-594, green), PRC1 (AF-647, magenta) and stained with DAPI (not shown). Images are sum-intensity projections of 5 z-planes smoothed with 0.5-pixel-Gaussian blur. Graph (right) shows MKLP1 intensity in untreated unlabeled (magenta) and BAC cells (green). The signal intensity of MKLP1 in these cells was 6% higher compared to endogenous MKLP1. (**E**) Time-lapse of HeLa BAC cell expressing MKLP1-GFP, depleted for endogenous PRC1, with transient expression of opto-PRC1 and iLID-CAAX (only MKLP1-GFP is shown) in time-points as in **Fig. 5 B** (top). Graphs (middle) correspond to membrane-to-membrane intensity profiles of MKLP1-GFP (green) and opto-PRC1 (magenta) retrieved from the same position perpendicular to the midzone in time-points as in **Fig. 5 B**, acquired by 1-µm wide line on maximum projection of three z-planes. Schematic representation of performed measurement is given on the right. Graphs (bottom) show individual MKLP1-GFP (left) and opto-PRC1 intensities. (**F**) Time-lapse of a cell as in **E**, which progresses to cytokinesis during time-points as in **E**. (**G**) Time-lapse images of U2OS cells with transient expression of opto-PRC1 (magenta) and GFP-Kif4A (green). Top (merge tgRFPt and SiR-DNA), bottom (GFP). Anaphase onset is set as time 0 min. (**H**) Spindle in a fixed untreated unlabeled U2OS cell (top) and unlabeled U2OS cell, depleted for endogenous PRC1 and transiently expressing Kif4A-GFP (not shown) and opto-PRC1 (not shown) (middle), immunostained for Kif4A (AF-647, green) and stained with DAPI (blue). Images are sum-intensity projections of 5 z-planes smoothed with 0.5-pixel- Gaussian blur. Graph (bottom) shows Kif4A intensity in untreated (magenta) and cells with transient expression of Kif4A-GFP (green). The signal intensity of Kif4A in these cells was 30% higher compared to endogenous Kif4A. (**I**) Time-lapse of U2OS cell as in **Fig. 5 B** (only GFP-Kif4A channel) (top). Graphs (middle) correspond to membrane-to-membrane intensity profiles of GFP-Kif4A (green) and opto-PRC1 (magenta) retrieved from the same position perpendicular to the midzone and in time-points as in **Fig. 5 B**, acquired by 1-µm wide line on a single z-plane. Schematic representation of performed measurement is given on the right. Graphs (bottom) show individual GFP-Kif4A (left) and opto-PRC1 (right) intensities. (**J**) GFP-Kif4A intensity on chromosomes in opto (black) and control (gray) cells 4 min after anaphase onset. Anaphase onset is defined as a timeframe of sister chromatid separation. Numbers in brackets; number of cells. Scale bars; 2 µm.

**Figure S6.**
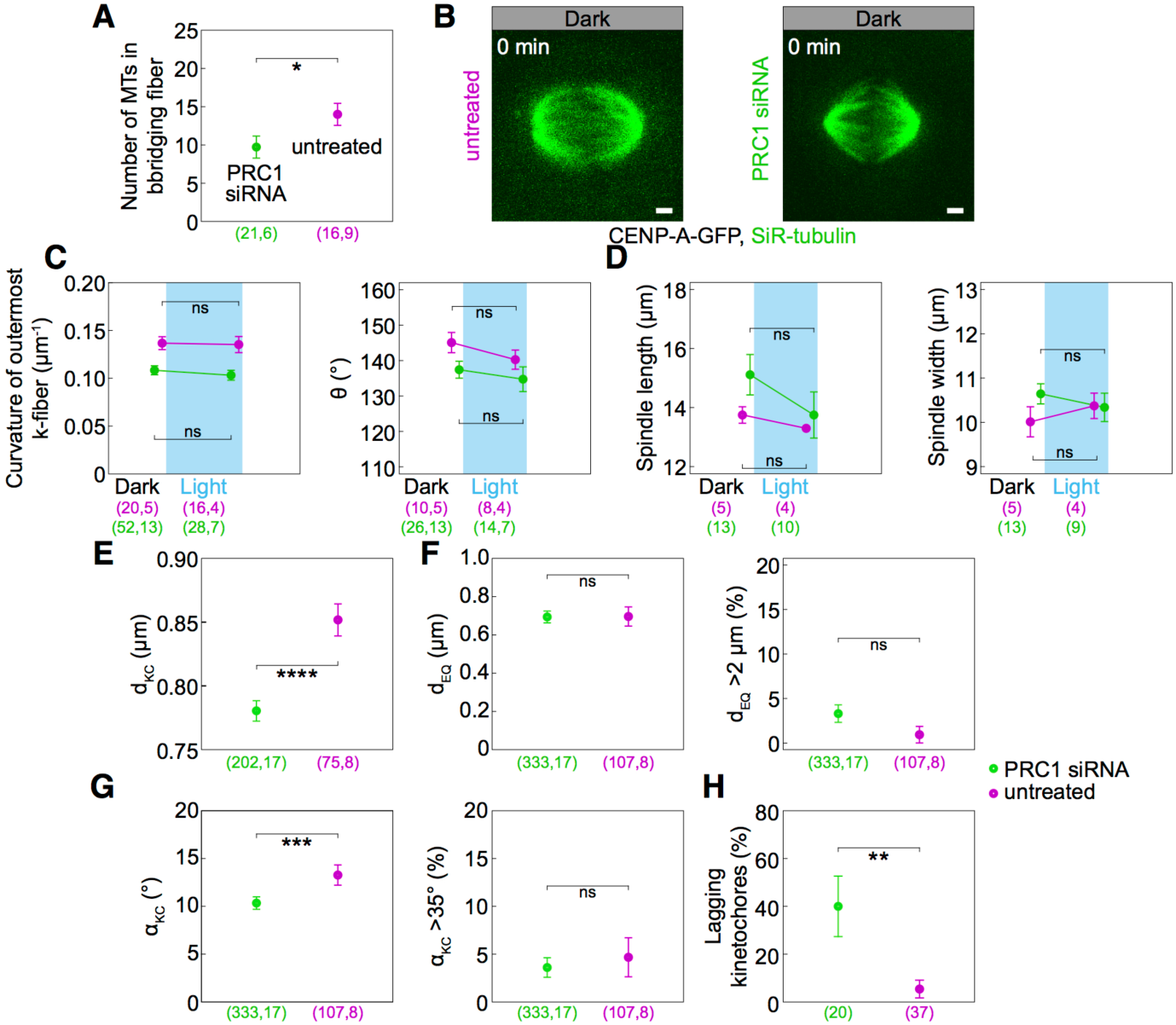
(**A**) Number of microtubules in the bridging fiber in untreated (magenta) and PRC1 siRNA treated HeLa cells (green) retrieved from our previous study (Polak et al., 2017). (**B**) Spindles of untreated (left) and PRC1 siRNA treated U2OS cells with stable expression of CENP-A-GFP (not shown) and stained with SiR-tubulin (green). (**C**) Curvature (left) and angle between (right) the outermost k-fibers in cells as in **B**, untreated (magenta) or PRC1 siRNA treated (green), before (0 min, Dark) and at the end of continuous exposure to the blue light (20 min, Light). (**D**) Spindle length (left) and width (right) in cells as in **B**. (**E**) Measurements of d_KC_ in untreated (magenta) and PRC1 siRNA treated (green) U2OS cells with stable expression of CENP-A-GFP and stained with SiR-tubulin. (**F**) Measurements of d_EQ_ in cells as in **B** (left) and percentage of d_EQ_ greater than 2 µm in untreated (magenta) and PRC1 siRNA treated (green) cells. (**G**) Measurements of α_KC_ in cells as in **B** (left) and percentage of α_KC_ greater than 35° in untreated (magenta) and PRC1 siRNA treated (green) cells. (**H**) Occurrence of lagging kinetochores in anaphase of untreated (magenta) or PRC1 siRNA treated (green) U2OS cells with stable expression of CENP-A-GFP and stained with SiR-tubulin. Cyan rectangles inside graphs indicate exposure to the blue light. Numbers in brackets denote measurements and cells; single numbers denote cells. Scale bars: 2 µm. Statistical analysis; t-test (**A**, **C-E**), Man Whitney test (**F** and **G**, left), two-proportions z-test (**F** and **G**, right; **H**); p-value legend as in **Fig. 2**.

## Video Captions

**Video 1.** U2OS cell stably expressing CENP-A-GFP (cyan), with transient expression of opto-PRC1 (magenta) and iLID-CAAX. The time of exposure to the blue light is indicated in the upper right corner. Time; min:s. The video corresponds to still images from **Fig. 1 B**.

**Video 2.** U2OS cell stably expressing CENP-A-GFP (cyan), with transient expression of opto-PRC1 (magenta) and iLID-CAAX. Note the kinetochores moving further from the equatorial plane upon PRC1 removal. The time of exposure to the blue light is indicated in the upper right corner. Time; min:s. The video corresponds to still images from **Fig. 2 B**.

**Video 3.** U2OS cell stably expressing CENP-A-GFP (cyan), with transient expression of opto-PRC1 (magenta) and iLID-CAAX, and microtubules stained with SiR-tubulin (green). Note the kinetochores lagging in the spindle midzone after anaphase onset. The time of exposure to the blue light is indicated in the upper right corner. Time; min. The video corresponds to still images from **Fig. 2 I**.

**Video 4.** U2OS cell stably expressing 2x-GFP-EB3 (grey) and mCherry-CENP-A (magenta), depleted for endogenous PRC1, with transient expression of opto-PRC1 (magenta) (top), and opto-PRC1 (magenta) and iLID-CAAX (bottom). The time of exposure to the blue light is indicated in the upper right corner. Time; sec. The video corresponds to still images from **Fig. 3 B**.

**Video 5**. U2OS cell stably expressing CENP-A-GFP (not shown), with transient expression of opto-PRC1 (magenta) and iLID-CAAX, and microtubules stained with SiR-tubulin (green).

The time of exposure to the blue light is indicated in the upper right corner. Top: SiR-tubulin; bottom: merge. Time; min. The video corresponds to still images from **Fig. 4 B** and **Fig. S4 A**.

